# The genetic control of rapid genome content divergence in *Arabidopsis thaliana*

**DOI:** 10.1101/2025.06.11.659220

**Authors:** Christopher J. Fiscus, Daniel Koenig

**Affiliations:** Department of Botany and Plant Sciences, University of California, Riverside, California, United States of America; Institute for Integrative Genome Biology, University of California, Riverside, California, United States of America

## Abstract

Genome evolution in eukaryotes is predominantly driven by the dynamics of repetitive sequences, which vary widely in both copy number and sequence composition. Rates of repeat evolution differ between and within species and are likely modulated by both genetics and environment. To uncover factors shaping the rate of genome content evolution, we analyzed 1,142 resequenced *Arabidopsis thaliana* genomes using a novel K-mer based approach to characterize genome content variation and identify hypervariable regions underlying differences in repeat abundance. We next treated repeat abundance as a quantitative trait and performed genome-wide association analyses across more than 400 repeat families to identify the genetic basis of copy number variation. Integrating these results through a meta-GWAS approach revealed both cis-acting variants and more than 50 trans-acting loci that regulate repeat abundance genome-wide. Cis-acting variation was predominantly localized to pericentromeric and centromeric regions, whereas trans-acting loci were enriched for candidate genes involved in DNA replication, DNA repair, DNA methylation regulation. Finally, we found evidence that purifying selection acts against mutations that accelerate genome content divergence, favoring alleles that constrain repeat expansion. Together, these findings provide new insights into the genetic architecture and evolutionary forces shaping genome evolution in *A. thaliana* and establish a framework for investigating these processes in other plant species.

## Introduction

Although core biological processes have been largely conserved across the tree of life, genome content and size vary dramatically, even between closely related species (Gregory, 2001). For instance, in eukaryotes, the number of genes per genome ranges from a few (Corradi et al., 2010) to tens of thousands, and genome size ranges several orders of magnitude (Elliott and Gregory, 2015). The lack of correlation between genome size and organismal complexity, called the “C-value enigma” (Gregory, 2001), was largely resolved by the discovery that eukaryotic genomes exhibit substantial interspecific variation in non-coding DNA (Stephan, 1989). Measurement of genome content by DNA hybridization (Britten et al., 1974; Britten and Kohne, 1968), followed by the generation of high-quality genome assemblies (Adams et al., 2000; Arabidopsis Genome Initiative, 2000; Lander et al., 2001), revealed that repetitive sequences comprise upwards of 90% of some eukaryotic genomes (Mehrotra and Goyal, 2014). Originally thought to be non-functional junk (Ohno, 1972), it is now generally recognized that repetitive sequences can be functional. For instance, telomeric repeats protect chromosome ends from degradation (Chakravarti et al., 2021) and centromeric satellite repeats play a role in kinetochore assembly during meiosis and mitosis (Talbert and Henikoff, 2022). Repetitive sequences can also have profound effects on phenotype by directly modifying gene expression and alternating local genetic and epigenetic mutation rates (Le et al., 2015; Slotkin and Martienssen, 2007). Beyond effects on phenotype, variation in repetitive sequence appears to be a major driver of genome content and size variation both over long (Novák et al., 2020) and short evolutionary timescales (Bosco et al., 2007).

The model plant *Arabidopsis thaliana* has a compact ∼150 Mbp genome (Davison et al., 2007) with over 10% variation in genome content between accessions (Long et al., 2013). Compared to its closest ancestor *A. lyrata*, from which it diverged < 6 million years (Hohmann et al., 2015; Novikova et al., 2016), the *A. thaliana* genome is repeat poor, with the majority of repetitive sequences occupying the centromere and pericentromere (de la Chaux et al., 2012; Hu et al., 2011). Population-level surveys of structural variation (Göktay et al., 2020; Lian et al., 2024), as well as repeat (Baduel et al., 2021; Davison et al., 2007; Lian et al., 2024; Long et al., 2013; Quadrana et al., 2016) and gene copy number variation (Lian et al., 2024; Zmienko et al., 2020), have previously been conducted in this system. Copy number variation and large indels in *A. thaliana* cluster in the repetitive sequences, often in the pericentromere, and more sequence losses have been observed than gains (Long et al., 2013). The majority of CNVs and indels overlap with transposable elements which are clustered in these regions. However, total TE content is relatively similar across sequenced accessions (Baduel et al., 2021). Instead the copy number of highly repetitive sequences in rDNA clusters (e.g. 45S rDNA) and in the centromeres are reported to make up a substantial portion of intraspecific variation in genome size and content (Long et al., 2013).

Efforts to catalog sequence differences segregating between individuals have historically relied upon variant calling and coverage-based approaches in which high-throughput sequencing reads from diverse individuals are mapped to a single reference genome. The strength of variant calling pipelines is their ability to detect small changes in conserved sequences, but variation in sequences absent from the reference genome are inaccessible to this method. For example, a study of genome variation in *A. thaliana* identified sequences up to 9 Mbp in length present in individual genomes but missing from the *Col-0* reference genome (Long et al., 2013). Alternatively, coverage-based approaches have been used to estimate copy number variation of genes and repeats. Repetitive sequences are largely missing or collapsed into single sequences in many first generation reference genomes (Treangen and Salzberg, 2011), and may be found in various sequencing contexts, hindering their detection by coverage differences. While recent advances in sequencing and assembly have ushered us into an era of “telomere-to-telomere” genome assemblies (Nurk et al., 2022) and bountiful pan-genomes (Cochetel et al., 2023; Golicz et al., 2016; Gordon et al., 2017; Lian et al., 2024; Liu et al., 2020; Montenegro et al., 2017; Zhao et al., 2018), the production and comparison of large numbers of genome assemblies remains cost prohibitive and technically challenging for many species. As such, the genomics community is investing substantially in developing methods to simultaneously compare very large numbers of eukaryotic genomes (Armstrong et al., 2020; Song et al., 2022; Zhou et al., 2024).

To complement variant calling and coverage-based approaches, we developed a K-mer based method to detect differences in genome content between individual genomes. K-mers are short sequences of length K that can be rapidly identified and counted in sequencing reads without the need for a genome assembly or alignment. K-mer profiles have been regularly used to compare and group genomes (Sievers et al., 2017). For instance, differences in K-mer profile have been used successfully to compare genomes in other systems such as maize (Liu et al., 2017) and *Drosophila* (Wei et al., 2014). Additionally, K-mer content has been used to estimate telomere length (Choi et al., 2021), estimate genome size variation (Sun et al., 2018) and as genotypes in GWAS in *A. thaliana* (Voichek and Weigel, 2020). Here, we first describe the development of our method before using it to catalog genome content variation in over 1,000 sequenced accessions of *A. thaliana*, focusing on copy number variation of repetitive sequences. We then mapped the genetic basis of repeat copy number variation for individual repeat families and used a “meta-GWAS” approach to identify trans-acting alleles affecting genome content variation more generally. Finally, we explore the evolutionary dynamics of associated variants and provide evidence that these variants appear to be under selection.

## Results

### Characterizing genome content using K-mer abundances from high-throughput sequencing data

We set out to characterize genomic variation in *A. thaliana*, with the goal of understanding which sequences vary in copy number between individuals. For this purpose, we explored using K-mer abundances in short read datasets to describe the content of a sampled genome, which we termed a genome content profile (GCP). We then used the genome content profiles to directly compare samples and to derive sequence copy number estimates for specific sequences of interest.

The first step in developing the method was to decide on the length of K-mer to use when generating the GCPs. Higher values of K make it easier to match sequences to their origin, but each increase in K grows the size of the dataset exponentially. As such, our priority was to identify the minimum value of K that could discriminate abundances of specific genomic sequences.

We started by comparing K-mer spectra in both repetitive and non-repetitive sequences in the *A. thaliana* reference genome for all values of K from 5 to 20. We reasoned that K-mers derived from repetitive sequences should be observed at a higher abundance than K-mers derived from non-repetitive sequences, if the length of K-mer was sufficiently large to discriminate between the two sequence types. We assessed when this condition was met by calculating the median abundance among the K-mers observed in each sequence bin (Fig. 1A). The minimum value of K required to discriminate the increased abundance of repetitive sequences was 12.

**Figure 1.**
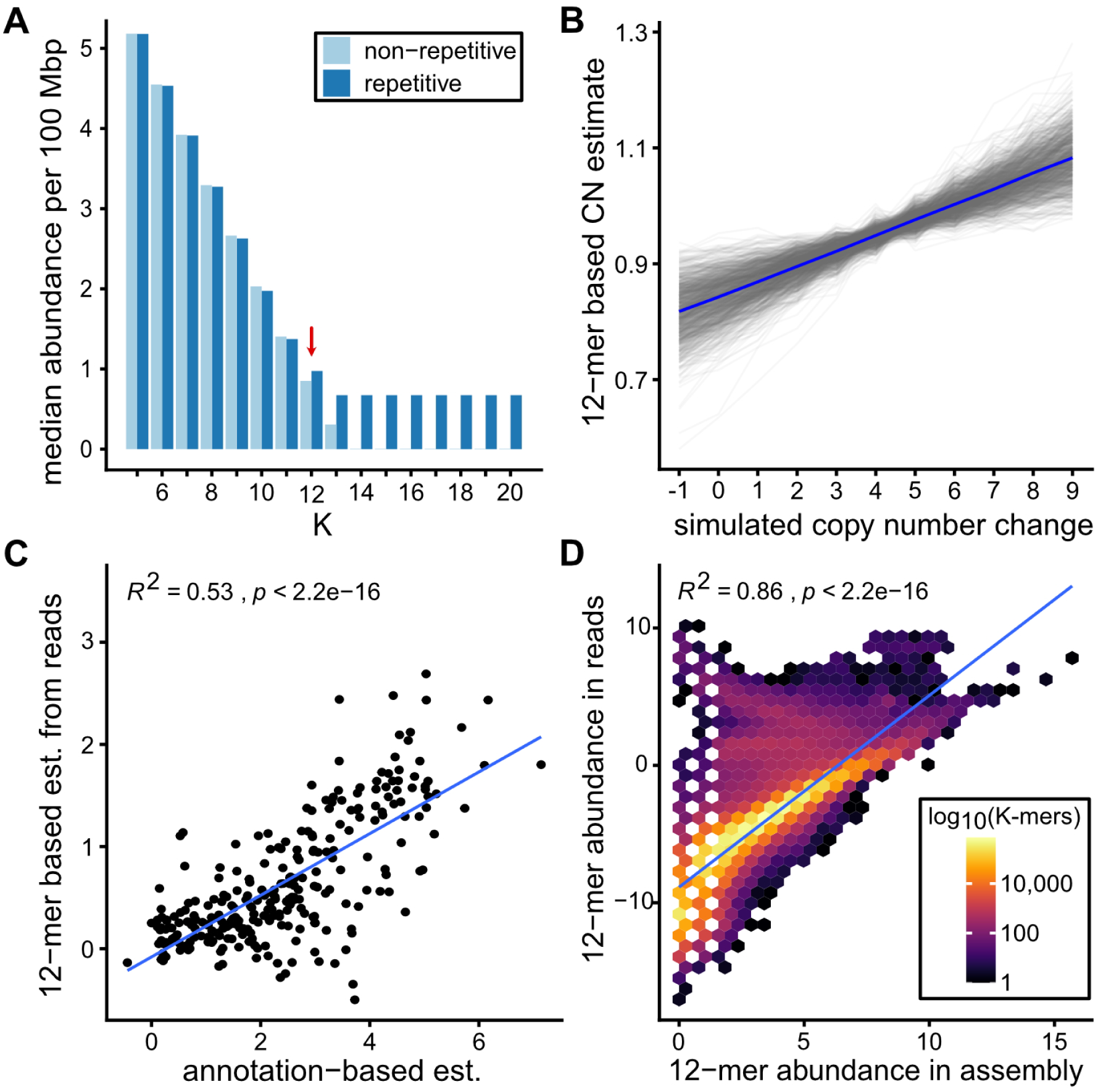
K-mer analyses of the *A. thaliana* reference genome. (A) Median K-mer frequency in repetitive and non-repetitive compartments of the reference genome. (B) Relationship between scaled 12-mer based copy number estimates (y-axis) and simulated copy number change across 1,000 simulations. Each gray line denotes a single simulation iteration, and the blue line indicates the median values across all iterations. (C) Relationship between 12-mer-based copy number estimates and annotation-based copy number estimates for 272 transposon families in the TAIR10 reference genome. The blue line indicates the best-fit line from a linear regression model. (D) Relationship between 12-mer abundances in Illumina reads (44 × mean coverage) and corresponding abundances in the genome assembly. Each hexagonal bin is colored by the number of observations within the bin. The blue line indicates the best-fit line from a linear regression model.

We next explored whether changes in sequence copy number in the *A. thaliana* genome might be detected using 12-mer abundances. We simulated copy number variation of 1 kb sequences in the reference genome and compared the known copy number with 12-mer copy number estimates. The median R^2^ between the simulated copy number changes and 12-mer based copy number estimates among 1,000 replicates was 0.98 (Fig. 1B, Fig. S1). We further validated this approach using published transposon abundances in the *A. thaliana* reference genome assembly (Cheng et al., 2017). We found 12-mer and annotation-based copy number estimates to be significantly associated in this dataset (Fig. 1C, R^2^ = 0.53, p < 2.2 X10^-16^).

Finally, we validated that 12-mer abundances in real and simulated Illumina sequencing reads (∼44X average genomic coverage) were strongly correlated with those in the reference genome assembly (Fig. 1D, R^2^ = 0.86, Fig. S2). However, the correlation between Illumina derived and simulated K-mer abundances are reduced when Illumina reads are downsampled to coverages less than 5X and break down at coverages less than 1X (Fig. S3). Thus, we concluded that moderate coverage whole genome sequencing datasets are appropriate for our method.

### A pipeline to generate genome content profiles for hundreds of genomes

To apply our approach to characterize genomic diversity in *A. thaliana*, we built a pipeline to assay genome content using K-mer abundances derived from large high-throughput sequencing datasets (Fig. S4). Raw sequencing reads were first processed and filtered to remove sequencing adapters, poor quality reads, and to correct likely base call errors. The filtered and processed reads were then mapped to the organellar genomes to filter reads likely originating from these sequences. 12-mer abundances were then counted in the unmapped reads. Finally, the 12-mer abundances for each sample were normalized for both GC content and coverage differences across samples. The normalized 12-mer counts were then used to estimate the abundance of sequences of interest for each sample based on the median abundance of constitutive 12-mers for that sequence (Materials and Methods).

We then applied our pipeline to profile the genomes of 1,319 resequenced *A. thaliana* collected from across the extant species range (1001 Genomes Consortium, 2016; Durvasula et al., 2017; Zou et al., 2017) (Table S1). To ensure that the analyzed dataset was high quality, we filtered samples that had low mappability, low coverage, extremes in %GC, or strong correlation between %GC and 12-mer abundance (Fig. S5). We further ensured that our dataset reflected the genomic variation in *A. thaliana* with minimal redundancy by clustering the individuals by genetic similarity and retaining one individual from each similarity group (static tree cut threshold of 100,000 corresponded to differences at ∼0.4% sites) for subsequent analysis. In addition to normalization to reduce the variation in GC content and coverage between samples (Fig. S6-S7), we fit a linear model to control for the effect of sequencing center bias, which we identified as a major factor driving variation in 12-mer abundance between individuals (Fig. S8). The final dataset represented 1,043 individual *A. thaliana* genomes with the normalized abundances of 8,390,656 discrete 12-mers. We hence refer to the normalized 12-mer abundances for each sample as a genome content profile (GCP).

### Regional copy number variation across the *A. thaliana* genome

One feature of the GCPs is that they are inherently reference-free. Therefore, to determine which regions of the *A. thaliana* genome contain the types of sequences that are variable between accessions, we segmented the Col-PEK genome assembly (Hou et al., 2022) into 100 kb non-overlapping windows and estimated the copy number variability of each window using the GCPs. For highly repetitive sequences this does not necessarily mean that the specific copy number variation occurs within that window, but it does indicate where the repeats that are most variable genome-wide occur. Copy number variability is herein described using the standardized range statistic (sR), which is defined as the range divided by the median of sequence abundance across individuals.

Copy number variability per genomic window, as measured by sR, spanned over two orders of magnitude (Fig. 2A). In general, sR had a strong positive correlation with the proportion of repetitive sequence within a window (Spearman’s *ρ* = 0.80, p < 2.2 × 10^-16^, Fig. S9) and a strong negative correlation with the proportion of genic sequence within a window (Spearman’s *ρ* = -0.87, p < 2.2 × 10^-16^). Regions with the highest extremes of sR almost exclusively corresponded to known cytological features including the pericentromere and centromere on all five chromosomes, the nucleolar organizer regions (rDNA) on chromosomes 2 and 4, and the heterochromatic knobs on chromosomes 4 (hk4S) and 5 (hk5L). In addition to heterochromatic regions of the genome, highly variable regions included a cluster of pre-tRNA genes on chromosome 1, clusters of cysteine-rich repeat secretory genes on chromosomes 3 and 4, and the RPP5 NBS-LRR disease resistance gene cluster on chromosome 4. These data are consistent with previous observations that some of these gene clusters show copy number variation between *A. thaliana* accessions (Choi et al., 2016; Zmienko et al., 2020).

**Figure 2.**
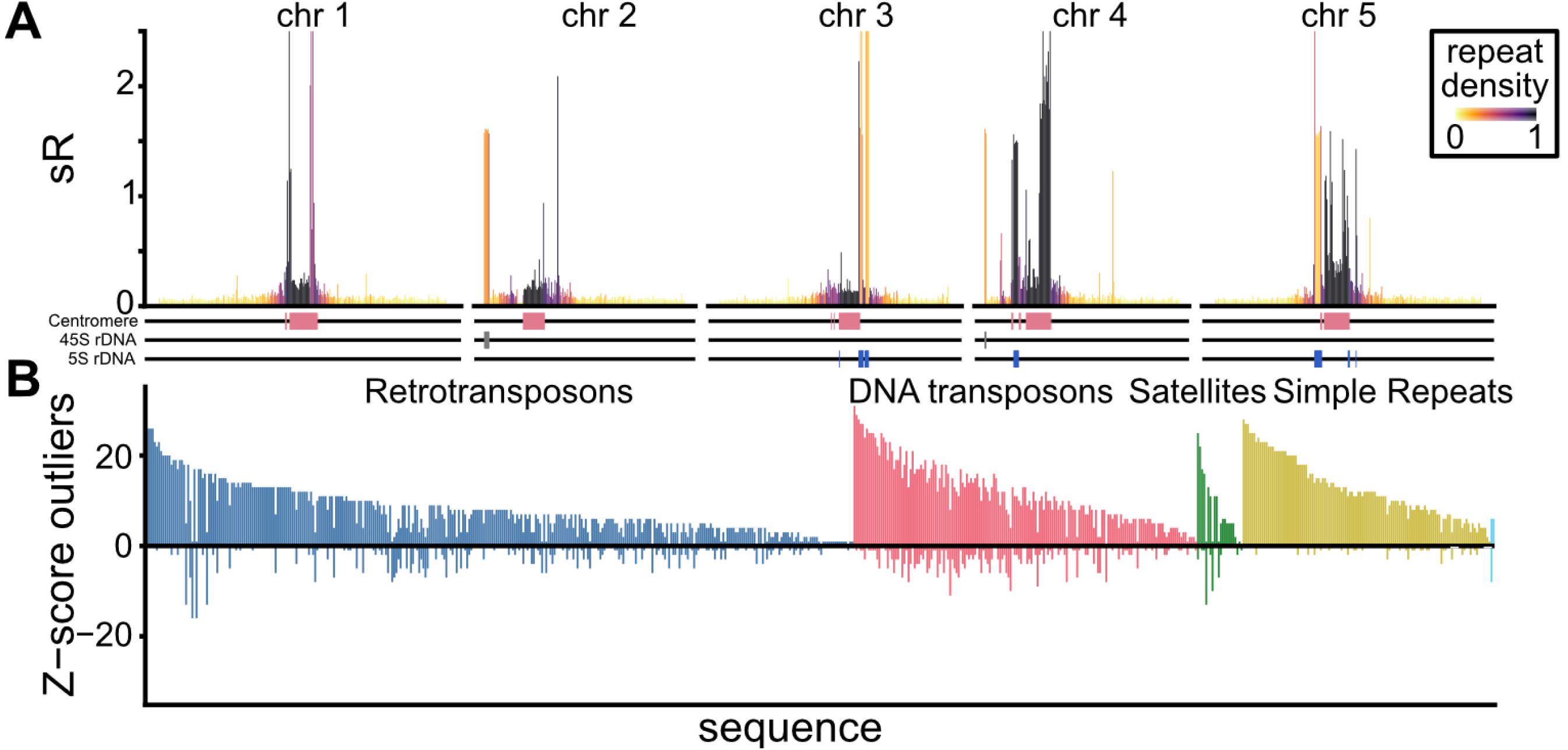
Patterns of copy number variability across the genome. (A) Sequence abundance variability in non-overlapping 100 kb windows across the genome in relation to repeat density and major cytological features. sR denotes the standardized range (range / median) of 12-mer abundance across all individuals. (B) Number of individuals with putative copy number changes per repetitive sequence. Z-scores were calculated from the 12-mer derived copy number estimates, and y-axis values represent the number of z-score values greater than 3 (copy number increase, positive value) or less than -3 (copy number decrease, negative value) across sequences. Sequences colored teal are unclassified repeats.

### Copy number variation of representative repetitive sequences

To determine which specific sequences vary between accessions, we next estimated the copy number of annotated sequences of interest in each genome using the GCPs (Table S2). We cataloged copy number variation of known repetitive sequences and of universal single copy orthologous genes (BUSCOs). We expected that BUSCOs would have lower rates of CNV than repetitive sequences based on their long-term intraspecific retention as single copy genes. Copy number variability per representative repeat sequence, as measured by sR, ranged nearly 4 orders of magnitude (0.07 to 500) compared to ∼1 order of magnitude for BUSCOs (0.08 to 0.69) (Fig. S10). On average, DNA transposons had higher sR than retrotransposons and satellites and simple repeats had higher sR than transposons (Wilcoxon-Mann Whitney U test, Bonferroni-adjusted p < 0.05).

We next identified repetitive sequences with strong changes in abundance over evolutionary time. We calculated a z-score for each repeat in each genome using the distribution of each repeat’s estimated abundance across accessions. Outlier z-scores (|z| > 3; p < .0027) represent repeats with unusually high or low estimated abundances in an accession (Fig. 2B). By this criteria, 98.6% of the tested repeats (648/657) changed in abundance in at least one accession, with an average of ∼11 accessions showing a change for any particular repeat (Table S2). Changes in repeat abundance were strongly enriched for expansions rather than deletions (6,050 increases and 914 decreases). Of the 657 representative repeat sequences, 348 exhibited only increases in abundance relative to the population mean. These included 195 retrotransposons, 52 DNA transposons, 12 satellite sequences, and 88 simple repeats, including all dinucleotide repeats. A total of 289 repeat families showed both increases and decreases in abundance, with a median ratio of accessions with increased versus decreased abundance of 3:1. This group comprised 137 retrotransposon sequences, 112 DNA transposon sequences (including all sequences from the mariner/Tc1 and pogo families), 8 satellite sequences, and 31 simple repeats. Only 11 repeat families exhibited decreases in abundance alone, including 8 retrotransposon sequences, the VANDALNX1 DNA transposon, and two satellite sequences (COLAR12 and the 5S rDNA repeat). An additional 11 repeat families showed no z-score outliers, including 7 Copia superfamily elements, the L1 superfamily element TA11, the DNA transposon VANDAL18, and two hexamer repeats (Fig. S11-S14).

### The evolution of repeat expansion and contraction in *A. thaliana*

We next investigated variation in repeat content across the 12 major subpopulations of *A. thaliana* previously defined using SNP data (1001 Genomes Consortium, 2016; Durvasula et al., 2017; Zou et al., 2017). Principal component analysis (PCA) revealed subtle shifts in repeat content across subpopulations along the first principal component (PC1; Fig. 3A). Loadings along PC1 indicated that these shifts were driven by copy number divergence across numerous repetitive sequences spanning all four major repeat classes (Fig. S15). Notably, DNA transposons showed particularly skewed loadings across multiple superfamilies, suggesting a disproportionate contribution to population-wide variation in repeat content (Fig. S16).

**Figure 3.**
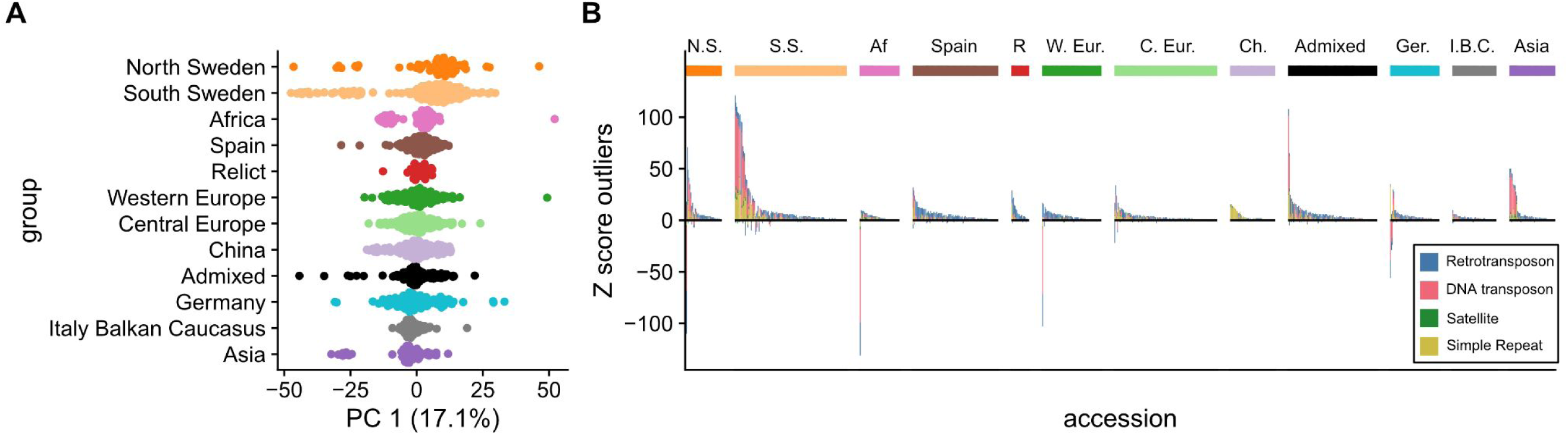
Repeat copy number differences between populations. (A) Principal component 1 of repeat copy number estimates partitioned by subpopulation. (B) Number of sequences with putative copy number change per individual grouped by subpopulation. Z-scores were calculated from 12-mer derived copy number estimates and y-axis values correspond to the count of z-score values greater than 3 (copy number increase, positive value) or less than -3 (copy number decrease, negative value) across individuals. Abbreviated subpopulations are as follows: N.S. is North Sweden, S.S. is South Sweden, Af. is Africa, R. is relict, W. Eur. is Western Europe, C. Eur. is Central Europe, Ch. is China, Ger. is Germany, I.B.C. is Italy Balkan Caucasus.

To assess whether repeat content reflects subpopulation structure, we tested for clustering by subpopulation. Repeat abundance estimates significantly clustered individuals in six of the twelve subpopulations–North Sweden, Italy Balkan Caucasus, Central Europe, Asia, Africa, and China–more often than expected by chance (permutation test, 10,000 permutations; Bonferroni-corrected p < 0.05). In contrast, we found no significant clustering for Western Europe, South Sweden, Germany, Relict, Admixed, or Spain. Despite limited clustering at the subpopulation level, the dominant pattern in the dataset was evidence of rapid, individual-level expansions and contractions of repetitive sequences (Fig. 3B). Consistent with this, the top 25 most divergent genomes were distributed across seven different subpopulations (Fig. S17).

Although global patterns of repeat content clustered significantly in only half of the subpopulations, we observed numerous instances of copy number changes that were significantly enriched within individual subpopulation groups. In total, 252 distinct sequences showed significant enrichment for copy number variation across 12 admixture groups (Fisher’s Exact test, Bonferroni-corrected p value across groups < 0.05, Table S3). South Sweden had the highest number of enriched sequences (N=138), followed by Asia (N=36) and Spain (N=22). In South Sweden, 135 sequences were enriched for copy number increases, including 78 DNA transposons, 17 retrotransposons, 4 satellites, and 36 simple repeats. This group also showed enrichment for copy number decreases in the 18S and 45S rDNA satellites and the ATMU8 MuDR superfamily DNA transposon. In Asia, copy number increases were enriched in 20 DNA transposons, 10 retrotransposons, 3 satellites, and 1 simple repeat, alongside a decrease in the ATCOPIA18A retrotransposon. Spain showed enrichment for copy number increases in 1 DNA transposon, 17 retrotransposons, and 2 simple repeats. The remaining groups had between 16 (Relict) and 2 (Admixed) sequences enriched for copy number change.

### The genetic basis of repeat copy number variation

Our analyses suggest that genome content variation primarily arises through changes in repeat-dense, heterochromatic compartments of the genome. Differences in repeat abundance between subpopulation groups are generally subtle, with a small subset of accessions distributed across groups exhibiting extreme genome content divergence. One possible explanation for this pattern is that these accessions may have accumulated one or more mutations that affect the likelihood of CNV occuring. To investigate this hypothesis, we treated the relative abundance of each repeat family as a quantitative trait and used a genome-wide association study (GWAS) approach to identify loci associated with repeat copy number. We tested 1,325,632 biallelic SNPs for association across 429 sequences, including 174 retrotransposons, 112 DNA transposons, all 22 satellites, and 121 simple repeats. In total, 29,891 SNPs were associated across 384 GWAS, including 142 retrotransposons, 58 DNA transposons, 21 satellites, and 58 simple repeats (Table S4).

### Pericentromeric regions are associated with repeat copy number variation

To identify genomic loci associated with repeat copy number variation, we first examined the genomic distribution of significant SNPs across all GWAS results (Fig. 4A). Significant associations were broadly distributed throughout the genome, with 94% of 100 kb windows (1,122 of 1,193) containing at least one significant SNP, and a median of 13 significant SNPs per window. This genome-wide pattern of significant SNPs was largely consistent across repeat classes. Despite the broad distribution, significant SNPs were enriched in pericentromeric regions, occurring more frequently than expected by chance (Fisher’s Exact Test, odds ratio 4.508, p < 2.2 × 10^-16^, Fig. 4B).

**Figure 4.**
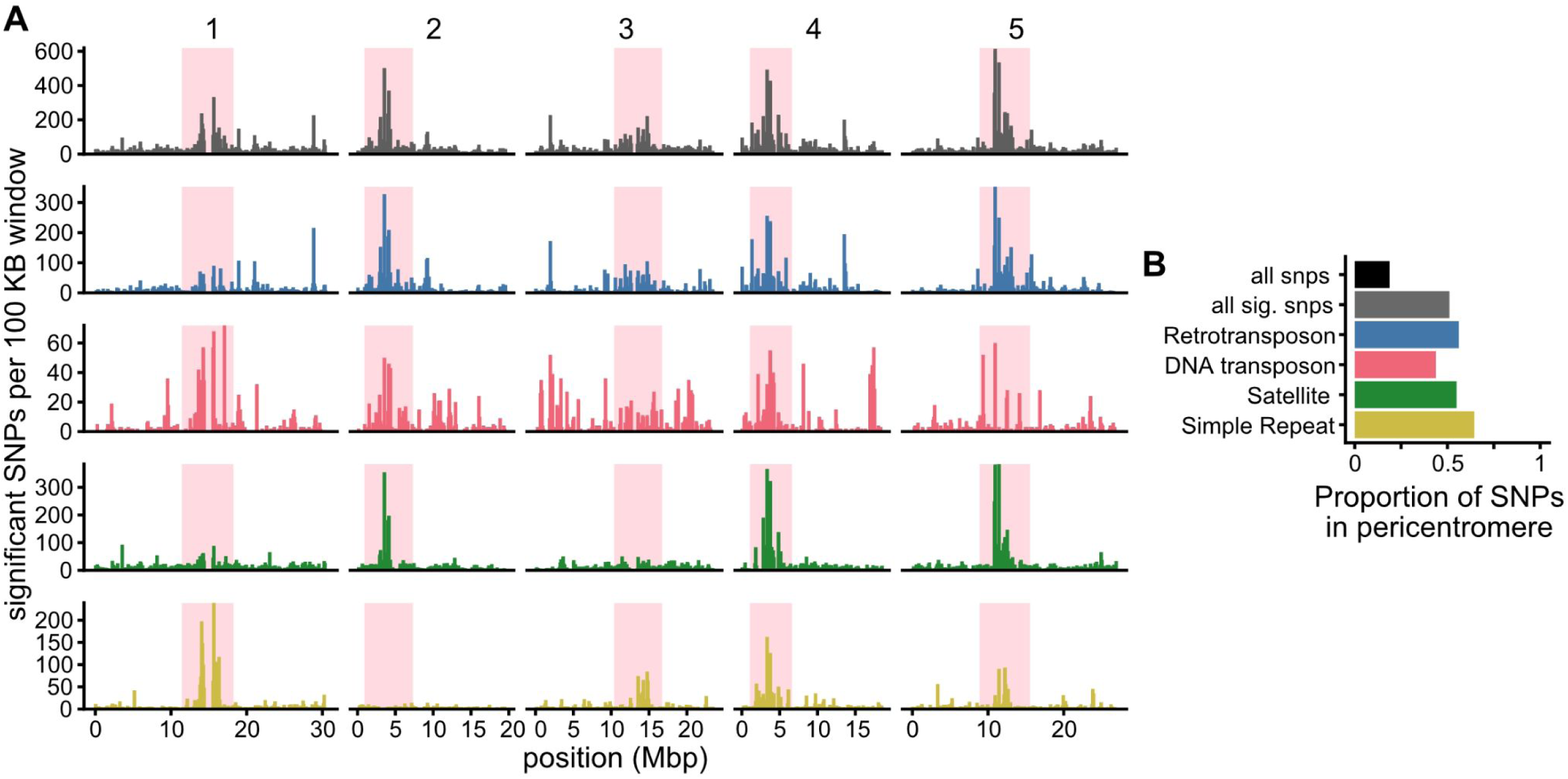
Genomic regions associated with repeat copy number variation. (A) Frequency of significant SNPs per 100 kb windows across all GWAS (gray), or GWAS in retrotransposons (blue), DNA transposons (red), satellites (green), and simple repeats (yellow), respectively. Pink highlighted region indicates centromeric and pericentromeric regions. (B) Proportion of all SNPs (black), significant SNPs from all GWAS (gray), and significant SNPs across GWAS by sequence class in the pericentromere.

Significant associations detected in our analysis may reflect either SNPs linked to copy number variants themselves (which we refer to as *cis*-acting associations) or SNPs linked to mutations that alter the rate that CNVs arise (i.e., putative *trans* mutators). Since repetitive sequences are concentrated within the pericentromeric and centromeric regions of *A. thaliana* (Fig. 2A, (Quesneville, 2020)), we hypothesized that the observed enrichment of significant SNPs in these regions likely reflects *cis*-acting variation in repeat copy number. To test this, we identified GWAS with enrichment for significant SNPs in the pericentromere, restricting our analysis to the 291 repeat families with at least 10 significant SNPs. We found significant pericentromeric enrichment in GWAS for copy number variation in 76 of 138 retrotransposons, 19 of 90 DNA transposons, 15 of 21 satellites, and 14 of 42 simple repeats (Fisher’s Exact Test, Bonferroni-corrected p < 0.05, Table S2). Of the retrotransposons tested, 45% (35 of 77) of Copia superfamily elements and 75% (40 of 53) of Gypsy superfamily elements showed significant enrichment of SNPs in the pericentromere. Similarly, all DNA transposons from the En/Spm superfamily, except ATENSPM1, were enriched in the pericentromere, along with ATHAT1 of the hAT superfamily, 41% (11 of 27) MuDR elements and 43% (3 of 7) VANDAL superfamily elements. In contrast, we detected no pericentromeric enrichment in GWAS for retrotransposons of the L1 or SADHU superfamilies or DNA transposons from the Helitron, mariner/Tc1, and pogo superfamilies. All satellites save for rDNA repeats and ATCLUST1 were enriched for centromeric SNPs. The GATCGATCGATC tetramer and some penta- and hexa-mers were also enriched for significant SNPs in the pericentromere. Supporting the interpretation that these associations reflect *cis*-linked CNVs, repeats with pericentromeric enrichment in GWAS were significantly more common in pericentromeric regions than in chromosome arms (Fisher’s Exact Test, odds ratio 10.349, p < 2.2e-16, Fig. S18).

Changes in pericentromeric repetitive sequences were not distributed evenly across the five pericentromeres (Table S5). In general, for transposons, there was a mixture of Copia and Gypsy superfamily LTR retrotransposons localized to all five pericentromeres, VANDAL superfamily DNA transposons localized to the pericentromeres on chromosomes 1-3, MuDR superfamily DNA transposons localized to the pericentromeres in chromosomes 2-5, and En/Spm superfamily DNA transposons localized to the pericentromeres on chromosomes 1-2. Notable sequences with sequence abundance variation localized to pericentromere 1 included a number of Copia transposons including *ONSEN* (ATCOPIA78), Gypsy transposons of the ATHILA8A family, satellite ATREP18 which contains the major telomere repeat, and centromere satellite variants from clusters 5 and 6. We localized abundance variation in AtSB5, a SINE non-LTR retrotransposon, and ATMSAT1, a mini-satellite, to the pericentromere on chromosome 2. Of note localized to pericentromere on chromosome 3 were all families of autonomous ARNOLD MuDR superfamily DNA transposons, and centromere satellite variant AR3. Pericentromere 4 was marked with variation in abundance of ATHAT1, the only hAT superfamily DNA transposon localized to a pericentromere, and numerous centromere satellite variants (AT12, COLAR12, cluster 2, cluster 3), as well as the major knob repeat of hk4S (ATENSAT1). Aside from transposons described previously, the only notable sequences localized to pericentromere 5 were variants of the hk5L knob repeat.

Taken together, our results suggest that much of the repeat copy number variation in *A. thaliana* can be attributed to genetic variation linked to pericentromeres.

### Trans-acting regulators of repeat copy number variation

We used two complementary approaches to identify *trans*-acting loci that may influence the abundance of multiple repeat families simultaneously. First, we conducted a meta-GWAS analysis by integrating association studies from 207 repeat sequences, including 104 retrotransposons, 34 DNA transposons, and 69 simple repeats. Second, we used the first principal component of repeat abundance variation as a quantitative trait in an additional genome-wide association study.

A large number of candidate *trans*-acting loci were revealed using the two approaches. The meta-GWAS yielded 228 significant SNPs, which collapsed into 57 discrete peaks after pruning for linkage disequilibrium (Fig. 5A, Table S6). GWAS on PC 1 yielded 84 significant SNPs, sharing 65 significant SNPs and 19 peaks with the meta-GWAS (Fig. 5B). The congruence of these analyses suggests that this approach captures loci generally associated with variation in repeat content between *A. thaliana* genomes.

**Figure 5.**
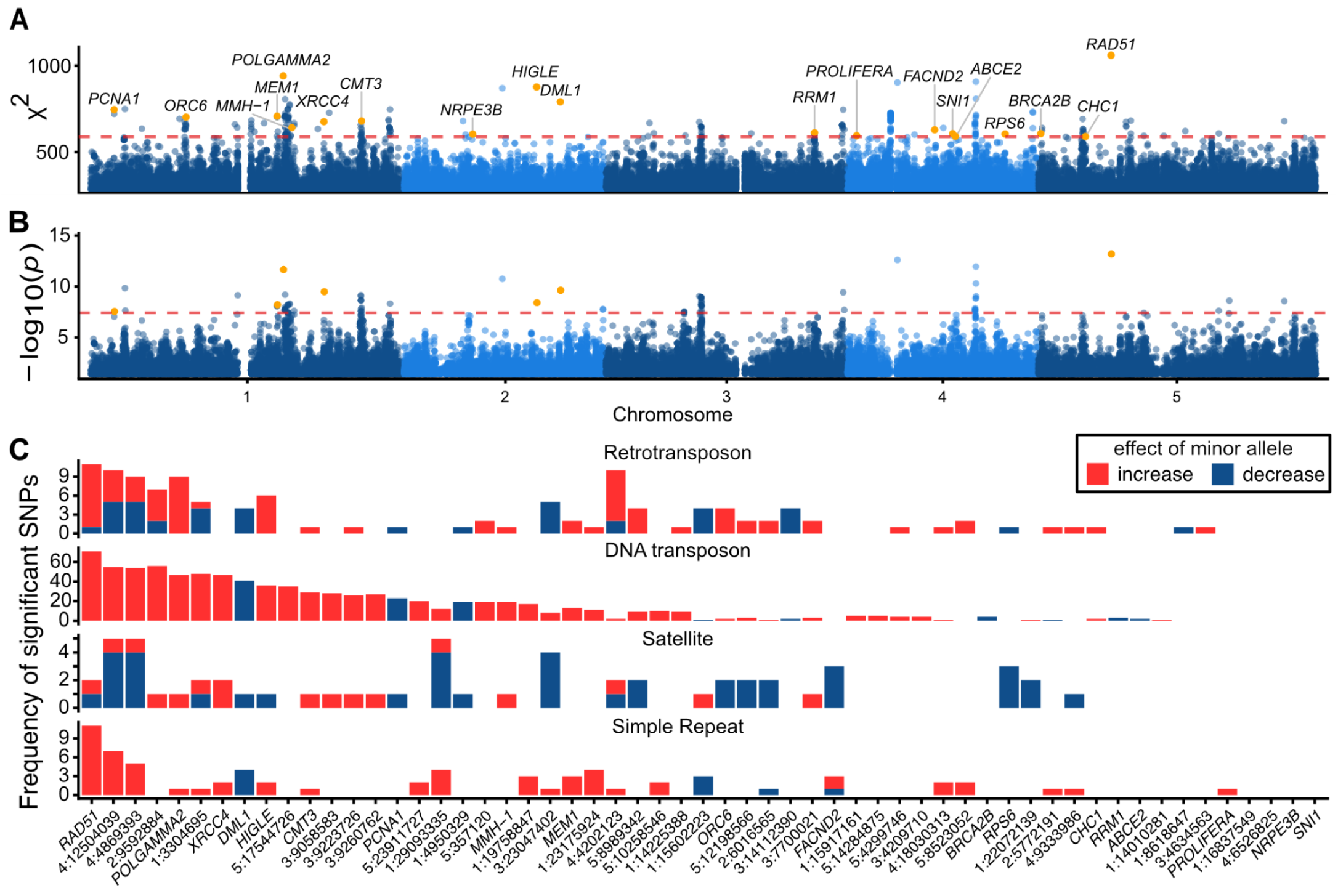
Genome-wide association mapping of repeat abundance. (A) Meta-GWAS of repeat abundance. The dotted line indicates the significance threshold at Bonferroni-corrected alpha = 0.05. Labels correspond to candidate genes at each locus listed in Table S6. (B) GWAS of PC 1 of repeat abundance. The dotted line indicates significance threshold at Bonferroni-corrected alpha 0.05. (C) Frequency of significant meta-GWAS tag SNPs across individual GWAS separated by sequence class. Bar colors indicate the effect of the minor allele on sequence abundance (i.e. positive beta corresponds to minor allele associated with increased sequence copy number).

Gene ontology enrichment analysis (GO analysis) for biological processes enriched in genes under meta-GWAS peaks yielded 63 significant GO terms at p < 0.05 (Table S7). Associated GO terms were involved in several biological processes including response to biotic and abiotic stresses (e.g. defense response to oomycetes, regulation of systemic acquired resistance, negative and positive regulation of plant-type hypersensitive response, and response to photooxidative stress), plant development (e.g. root development, floral whorl development, and maintenance of seed dormancy by abscisic acid), and DNA replication and repair (e.g. DNA repair, premeiotic DNA replication, and mitotic recombination-dependent replication fork processing). The latter of these enrichments suggested that changes near genes involved in DNA repair may be associated with genome divergence *A. thaliana*.

We subsequently manually examined the meta-GWAS peaks to identify potential candidate genes with activity linked to repeat copy number variation. Consistent with the results of the GO analysis, we found more than a dozen genes involved in DNA replication and/or DNA repair pathways underlying the peaks (Table S6). In total, we identified candidate loci for 21 of the peaks, with 9 tag SNPs predicted to be missense variants or linked to predicted missense variants (R^2^ > 0.20). We hereafter identify genes tagging putative missense variation with an “*”.

Candidate genes involved in DNA replication included *ORC6\** (AT1G26840), a subunit of the DNA replication origin recognition complex (Masuda et al., 2004), *PCNA1* (AT1G07370), a sliding clamp that acts in leading strand elongation (Strzalka and Ziemienowicz, 2011), and *PROLIFERA* (AT4G02060), a component of the minichromosome maintenance complex (Springer et al., 2000).

Candidate genes involved in DNA repair were primarily associated with the repair of DNA double strand breaks via homologous recombination. Examples include *HIGLE* (AT2G30350), a SLX1 family endonuclease that resolves Holliday junctions (Verma et al., 2022), *FANCD2\** (AT4G14970) a homolog of Fanconi anemia D2 that is essential for homologous recombination during meiosis (Kurzbauer et al., 2018), the homolog of *RAD51* (AT5G20850) (Doutriaux et al., 1998), involved in homology search during homologous recombination (Li et al., 2004), and the homolog of breast cancer susceptibility gene 2 (*BRCA2B*, AT5G01630) which recruits *RAD51* and *DMC1* during double-strand break repair via homologous recombination (Seeliger et al., 2012). Additional candidates involved in DNA double strand break repair included *XRCC4\** (AT1G61410), a homolog of a DNA ligase IV binding protein with a role in the non-homologous end joining pathway (West et al., 2000) and DNA polymerase gamma-2 (*POLGAMMA2*, AT1G50840), an error-prone organellar DNA polymerase encoded in the nucleus (Ayala-García et al., 2018).

Candidate genes implicated in the repair of other classes of DNA damage were also observed under meta-GWAS peaks. Examples include MutM homolog *MMH-1\** (AT1G52500), which is involved in the repair of single strand nicks due to oxidative DNA damage during base excision repair (Ohtsubo et al., 1998) and *CHC1* (AT5G14170), a subunit of SWI/SNF chromatin remodeling complex involved in repair of DNA damaged by UV-B (Campi et al., 2012). We also found *SNI1\** (AT4G18470), a subunit of the Smc5/6 complex that reacts to DNA damage by acting as a DNA micro-compaction machine that identifies unusual DNA conformations (Serrano et al., 2020).

In addition to genes involved in DNA repair, we found many candidates with roles in DNA methylation and DNA demethylation, which likely act in the epigenetic regulation of repetitive sequence proliferation. Notable candidates involved in DNA methylation included *CMT3* (AT1G69770), a plant-specific chromomethylase which methylates cytosine at non-CpG sites (Lindroth et al., 2001) and *RRM1\** (AT3G54760), which interacts with *SUVH9* to regulate RNA-directed DNA methylation (Zhou et al., 2021). Candidates acting in DNA demethylation included *DML1*/*ROS1\** (AT2G36490), which demethylates DNA (thus repressing gene silencing) (Gong et al., 2002) and *MEM1* (AT1G48950), which removes CHH methylation (Lu et al., 2020). Further candidates included *NRPE3B\** (AT2G15400), a subunit of plant-specific RNA polymerase V with a role in siRNA-directed DNA methylation (Ream et al., 2009), *ABCE2* (AT4G19210), a RNase L inhibitor that suppresses RNA silencing (Braz et al., 2004), and *RPS6* (AT4G31700) a component of the 40S ribosomal subunit that represses rDNA transcription (Kim et al., 2014).

We examined the extent that the 57 loci identified in the meta-GWAS were significant in individual GWAS for repeat copy number change. In total, 53 loci were significant in one or more GWAS, with 12 associated with GWAS in all four repeat classes, 16 found in three repeat classes, 14 found in 2 repeat classes, and 11 found in only one repeat class (Fig. 5C, Table S4). Four loci were significant in the meta-GWAS but were not significant in any individual sequence copy number GWAS nor in the PC 1 GWAS. The former loci included the peaks with candidates *SNI1* and *NRPE3B* as well as two loci without an identified candidate.

To understand whether associated peaks have consistent effects on different repeat classes we characterized the effect of the minor allele on sequence copy number among the individual GWAS for each peak. The majority of loci promote copy number variation across different repeats in the same direction (66%, 26 increasing and 9 decreasing abundance, Fig. 5C). Among the peaks with mixed effects, 5 peaks have mixed effects in two or more classes and 3 have mixed effects in one class only. These data indicate that most of the putative mutator alleles that we have identified in *A. thaliana* generate mutations that are more likely to increase copy number. How these loci are generating directional shifts in repeat copy number remains to be explored.

### Evolutionary dynamics of putative mutator alleles

Finally, we calculated the species-wide site frequency spectrum to assess whether loci associated with repeat copy number variation are subject to selection. We observed a significant enrichment for rare alleles (minor allele frequency < 0.10) among both meta-GWAS tag SNPs and SNPs significant in one or more GWAS, relative to nonsynonymous SNPs (Fig. 6A, Chi-squared tests, p = 0.031 and p < 2.2 × 10 ^-16^, respectively). However, there was no significant enrichment of rare SNPs in the meta-GWAS tag SNPs when compared to putatively causative SNPs from the AraGWAS database (Togninalli et al., 2020, 2018), even when stratifying meta-GWAS SNPs by the direction of derived allele effect. Our results suggest that variation associated with repeat copy number change is likely under purifying selection, as indicated by the excess of low-frequency alleles.

**Figure 6.**
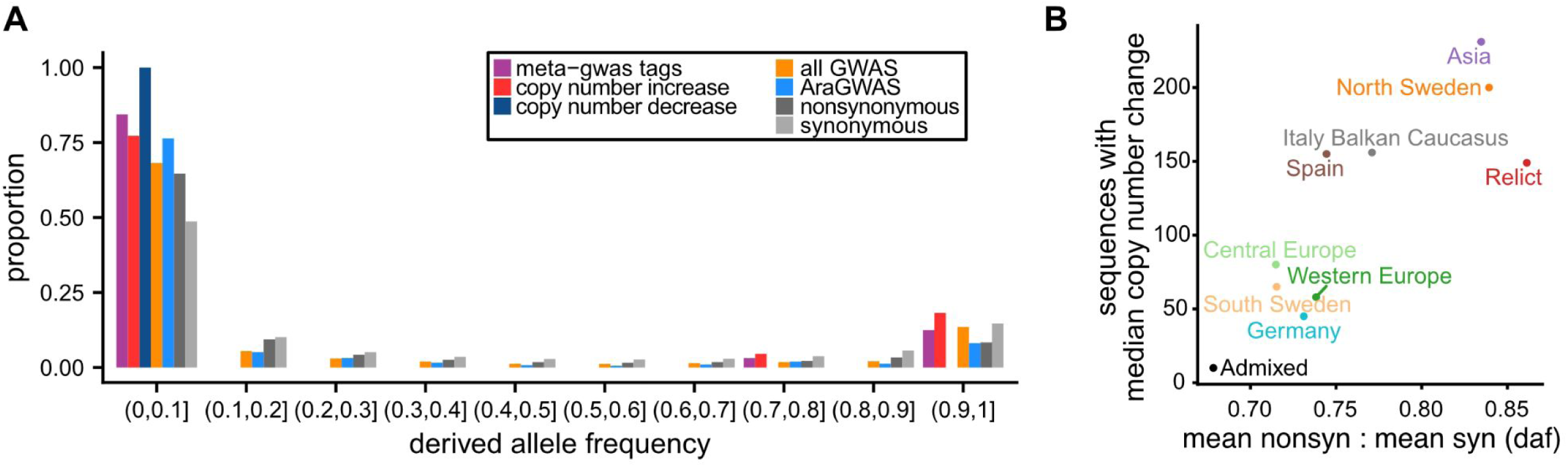
Evolutionary dynamics of alleles associated with copy number change. (A) Site frequency spectra of meta-GWAS tag SNPs, meta-GWAS tag SNPs stratified by predominant copy number effect, all GWAS-associated SNPs identified in this study, and AraGWAS-associated SNPs, compared to nonsynonymous and synonymous SNPs from the 1001 Genomes dataset. (B) Relationship between the median number of sequences with copy number change per subpopulation (normalized to the Africa group) and the ratio of mean derived allele frequencies between nonsynonymous and synonymous SNPs.

If purifying selection is acting to purge mutations that drive rapid genome content divergence, then the strength of selection within a population should influence the rate of genome evolution. We found that many of the accessions with evidence of rapid genomic divergence occur outside of the core of the species’ European range, where effective population sizes are smaller. To test whether repeat differences between populations could be attributed to variation in the strength of selection, we examined the relationship between the number of sequences with copy number changes per group and the strength of genetic drift, estimated by the ratio of derived allele frequencies between nonsynonymous and synonymous variants (Fig. 6B). We observed a strong positive correlation (Spearman’s *ρ* = 0.76, p = 0.02) between these parameters, suggesting that populations with more extensive copy number changes have likely experienced relaxed selection.

## Discussion

### A K-mer based method to estimate sequence copy number

Here, we present a novel K-mer based approach to estimate sequence copy number from high-throughput sequencing reads and apply the method to study genome content variation in a species-wide survey of wild *A. thaliana.* The method is designed to profile genomes sequenced with short read technology while avoiding the complexities of genome assembly. As such, the method is useful to study sequences that present difficulties for genome assembly such as repetitive sequences (Treangen and Salzberg, 2011), but it has the disadvantage of being unable to localize a sequence in a genome. While we have entered the era of complete (i.e. telomere-to-telomere) genome assemblies (Nurk et al., 2022), such assemblies take considerable resources to produce and will likely be infeasible for studying genomic variation at population scale for some time. In contrast, our method does not require high-coverage sequencing or long reads and thus represents a cost-effective method to study genomic variation in many samples. Our method can also be readily applicable to the extensive library of high-throughput sequencing reads in public databases such as NCBI SRA.

One important consideration when running the pipeline is the choice of K-mer length. We describe a method here to choose K and used K = 12 for our analyses, based on empirical observations of K-mer abundance in repetitive versus non-repetitive compartments of the reference genome (Fig. 1A). Still, we admit that this choice is somewhat arbitrary, and that considering multiple K-mer lengths is likely valuable. Longer K-mers offer better sequence specificity, but they carry a greater risk that mutations within the sequence will disrupt the K-mer and bias abundance estimates; shorter K-mers are less susceptible to this issue but likely lack the specificity to distinguish distinct repeat sequences. We ultimately erred on the side of choosing a smaller K, since each increase in K grows the dataset exponentially and inflates the number of uninformative singletons (i.e. K-mers with abundance 1). Because the choice of K = 12 was not obvious, we also produced GCPs using 10-mers and repeated the GWAS for a subset of sequences.

Comparing the two lengths showed that K can meaningfully change which loci are mapped, with neither length uniformly more powerful. The effect was clearest for three repeats. At one extreme, the 12-mer GWAS for *ONSEN*, the well-characterized heat-activated retrotransposon in *Arabidopsis* (Cavrak et al., 2014) produced a single peak in the pericentromere of chromosome 1, while the 10-mer GWAS yielded no significant hits at all (Fig. S19). The DNA transposon ATHATN2 showed the opposite pattern: here the 10-mer GWAS recovered more of the candidate SNPs identified by our meta-GWAS than the 12-mer version did (Fig. S20). The GWAS for VANDAL1, another DNA transposon, fell somewhere in between, with the two lengths implicating different regions: the 12-mer GWAS mapped variation to the pericentromeres of chromosomes 2, 3 and 5, while the 10-mer GWAS only mapped to the pericentromere of chromosome 2 (Fig. S21).

Ideally, we would have also generated GCPs using a larger K. We attempted this, but given our current pipeline design and methodology, we found the task to be impractical for the large number of genomes included in our analyses. Refactoring the pipeline to support data structures other than a single large hash table, such as a database or per-sample storage of K-mer counts, paired with filtering of singleton K-mers to reduce the dataset size, are possible future improvements to the method. At the same time, our results with the 10-mer profiles demonstrate that smaller K-mers have real value. Despite their reduced specificity, 10-mer based GWAS recovered some signal that the 12-mer GWAS missed, underscoring that larger K are not uniformly superior and that shorter K-mers merit continued consideration alongside efforts to scale the pipeline to handle larger K.

We used the genome profiles generated by our pipeline to estimate sequence copy number using the abundance of K-mers that constitute a focal sequence. This application is dependent on providing reference sequences from which the K-mer hash table is derived. For our work, we primarily focused on studying copy number variation in known repetitive sequences previously identified in *A. thaliana* and contained within RepBase (Bao et al., 2015). However, comprehensive repeat libraries are not yet available for the majority of systems. While approaches to construct such libraries do exist (e.g. EDTA (Ou et al., 2019)), contemporary approaches require high-quality genome assemblies with assembled repeats. Additionally, our method imperfectly accounts for variation in the sequence composition of representative sequences. Although the method can accommodate sequences containing IUPAC nucleotide ambiguity codes, the current implementation expands each ambiguous K-mer into all possible nucleotide combinations and substitutes the median abundance of these possible K-mers when calculating the abundance estimate. When multiple ambiguous nucleotides occur within a single K-mer, the number of possible combinations increases exponentially, reducing performance. Future implementations of the pipeline could incorporate a weighting scheme to better handle conserved and variable sequences within representative sequences when estimating repeat abundance.

A major issue with the approach is that K-mer profiles seem to be especially sensitive to technical biases in sequencing data, such as batch effects, that are typically unaccounted for in genomic analyses. We discovered this effect by doing a principal component analysis of normalized 12-mer profiles and realized that samples clustered according to the sequencing center (Fig. S8). We investigated this issue by attempting to isolate the variable responsible for the technical bias. Specifically, we examined GC content differences between libraries, possibility of organellar genome contamination, and the effect of low and high abundance 12-mers on genome content profiles. Ultimately, we were unable to determine the cause of the bias. In particular, within the 1001 Genomes dataset, population group and sequencing center were confounded, making it impossible to distinguish between biological and technical variation. To mitigate this issue, we fit a linear model to partially account for the effect of sequencing center on genome content profiles, with the understanding that we were also likely removing some biological variation. Future applications of this approach should incorporate experimental designs that enable robust estimation of technical effects, such as sequencing replicate samples randomized across sequencing runs, and apply statistical approaches to account for batch effects in high-dimensional genomic count data..

### Genome content variation in wild *A. thaliana*

Our results suggest that genome divergence between *A. thaliana* accessions is largely explained by copy number variation of repetitive sequences in heterochromatic-rich regions of the genome such as the nucleolar organizing regions, pericentromeres, and centromeres. This finding is consistent with very high collinearity observed in gene-rich regions of 69 chromosome-scale *A. thaliana* genomes (Lian et al., 2024). The implication is that highly repetitive sequences are probably among the most rapidly evolving sequences within the genome and therefore could underlie genomic changes that eventually lead to speciation. Consistent with this possibility, comparisons between *A. thaliana* and its closest related species *A. lyrata* found extensive divergence in repetitive DNA. The ∼70 Mbp difference in genome size between the two species is primarily attributed to the purging of non-coding DNA, including transposable elements, in the lineage leading to *A. thaliana* (Hu et al., 2011). Furthermore, comparisons of centromere assemblies between the two species showed complete divergence of the orthologous centromeres (Wlodzimierz et al., 2023). Extensive sequence divergence has also been found in copies of the 178 bp canonical major centromeric satellite repeat (*AthCEN178*) both within (Maheshwari et al., 2017) and between *A. thaliana* genomes, with almost a quarter of individual repeat copies unique to a specific genome (Wlodzimierz et al., 2023).

An alternative, non-exclusive explanation for this divergence pattern is that purifying selection is much weaker in regions of the genome that are repeat-rich (and thus gene-poor). This view is supported by evidence that TEs can insert all over the genome and seem to lack a preference for inserting within the repeat-rich pericentromeres and centromeres (Quadrana et al., 2016). Thus, TEs that insert near genes are rapidly purged by strong purifying selection (Hollister and Gaut, 2009), with selection pressure greater than that exerted on missense SNPs (Baduel et al., 2021). Mechanistically, the weakening of purifying selection in repeat rich regions could be a local reduction in recombination rate in repeat-rich regions of the genome. This is especially true of centromeres, which have been shown to have both low and non-recombining compartments (Fernandes et al., 2024).

Overall, we observed many more instances of repeat copy number increase compared to decrease. This pattern could be indicative of repeat expansions being more frequent than contractions or due to stronger purifying selection against copy number decreases compared to increases. Although our data does not allow us to discriminate between these two possibilities, the enrichment for rare variants was stronger for alleles associated with copy number decrease compared to increase (Fig. 6A), lending credence to the latter hypothesis. The latter hypothesis is further supported by the observation that most transposons in the *A. thaliana* genome appear to predate speciation (Quesneville, 2020), suggesting that deletions of these sequences would be strongly deleterious. It remains to be seen whether stronger selection against further deletions is the result of the already extensive genome reduction in *A. thaliana* making further deletion more likely to disrupt biological function.

Our work has many similarities with a smaller survey of transposon copy number variation in 211 accessions (Quadrana et al., 2016). Of the 174 transposon superfamilies analyzed in both studies, the presence or absence of copy number variation was consistent between the previous study and our work for 80 superfamilies. In contrast, we identified copy number variation in 30 superfamilies not identified in the smaller cohort while they detected CNV in 64 superfamilies which were not detected in our work. Despite these discrepancies, 24 of 59 comparable GWAS had one or more shared peaks between studies. It is unclear whether coverage-based or K-mer based copy number estimates generated from sequencing reads are more reliable indicators of transposon copy number variation between genomes. We anticipate that repeat analysis in chromosome-scale genome assemblies (Kang et al., 2023; Lian et al., 2024; Wlodzimierz et al., 2023) will clarify which families have copy number variation in *A. thaliana*.

### The genetic basis of repeat copy number variation

In addition to mapping *cis-*acting repeat copy number variants, we identified over 50 distinct *trans*-acting loci regulating repeat copy number variation. These *trans*-acting loci were enriched for genes involved in DNA repair, DNA replication, and DNA methylation pathways. Notably, among the candidate genes associated with DNA repair, the majority function in the repair of DNA double-strand breaks (DSBs), which is consistent with the role of DSBs in promoting copy number variation through mechanisms such as transposon activity (Turlan and Chandler, 2000) and recombination events that underlie the expansions of simple repeats and satellites (Gadgil et al., 2020).

We also identified candidate genes involved in other DNA repair pathways, including nucleotide excision repair, which resolves intrastrand crosslinks and bulky adducts, and base-excision repair, which repairs single strand lesions. While these latter pathways may not directly mediate repeat amplification, they can repair collateral DNA damage associated with the proliferation of repetitive elements. For example, in the cut-and-paste replication of DNA transposons, transposase activity can introduce single-strand breaks at the donor site (Hickman and Dyda, 2015). Additionally, the DNA repair processes themselves can generate sequence copy number variation: single strand breaks are extremely common in plant genomes (Wolter et al., 2021) and have been directly linked to the formation of tandem duplications of short repeats (Schiml et al., 2016).

Our work suggests that rapid changes in genome content arise sporadically throughout *A. thaliana*’s range, in part due to mutations that alter the efficiency of various DNA repair pathways and DNA methylation processes. Changes in genome content appear to be deleterious, as they are associated with rare genetic variants and occur more frequently in populations where purifying selection is less effective. We expect that both our ability to detect the effects of these mutations and the efficacy of selection in shaping their consequences are influenced by *A. thaliana*’s predominantly self fertilizing mating system. In outcrossing species, mutator loci are rapidly segregated away from the CNV that they induce. In contrast, a self fertilizing lineage that acquires one or more mutator alleles may be doomed to accumulating large numbers of deleterious CNVs unless rescued by an outcrossing event that allows mutators to be recombined away from their deleterious genomic consequences. Our ability to detect these lineages in *A. thaliana* suggest that they either occur at relatively high frequency, or persist for several generations.

Together, these data suggest a model in which mutator loci emerge in a self fertilizing lineage, induce elevated copy number variation rates, and persist when purifying selection is weak or the consequences of the specific genomic changes in content are small. If outcrossing occurs, progeny that do not carry the mutator loci could retain changes in genome content. Populations with small effective population size, where this cycle may occur more frequently, could be placed on a path of rapid genomic content evolution. This model may help explain the episodic bursts of repeat expansions that characterize genome content evolution across the plant kingdom.

## Materials and Methods

### Genome annotation

To identify repetitive and non-repetitive bins of the *A. thaliana* genome, we annotated the reference genome assembly (TAIR10) (Lamesch et al., 2012) with RepeatMasker 4.1.1 (Smit, AFA, Hubley, R & Green, P, 2013-2015) using the Dfam 3.2 library (Storer et al., 2021) and specifying “-species arabidopsis”. Presumed conserved single copy genes were identified by annotating the reference genome with BUSCO 3.02 (Simão et al., 2015; Waterhouse et al., 2018) using the Eudicotyledons OrthoDB release 10 (Kriventseva et al., 2019) library. Augustus 3.3 (Stanke et al., 2008) was used as a dependency for the BUSCO pipeline.

### K-mer analyses of TAIR10 reference genome

Canonical 5 - 20 mers were counted in repetitive and non-repetitive sequences of the *A. thaliana* reference genome using jellyfish 2.2.9 (Marçais and Kingsford, 2011). Designation of sequence as repetitive or non-repetitive was based on the RepeatMasker annotation described previously. From these K-mer counts, the number of unique K-mers and the median K-mer abundance per 100 Mbp bin were calculated.

To demonstrate that 12-mer profiles generated using the pipeline from sequencing reads reflect 12-mer profiles from genome assemblies, canonical 12-mers were calculated using jellyfish as described above for the reference genome and compared to normalized 12-mer counts produced by the software pipeline depicted in Fig. S4. A linear model was fit to log-transformed 12-mer counts using R (R Core Team, 2013).

To explore how coverage affects the relationship between the respective 12-mer profiles generated from sequencing reads and genome assemblies, Spearman’s correlation was calculated between 12-mer profiles generated from *Col-0* whole-genome sequencing reads (1001 Genomes Consortium, 2016) and the reference genome. For the sequencing reads, both empirical and simulated data were considered at coverages of 20, 10, 5, 2.5, 1, 0.5, 0.25, and 0.1X. Seqtk 1.2-r95 (Li, n.d.) was used for sampling of empirical data and wgsim 1.9 (Li et al., 2009) was used to simulate 100 bp reads from the TAIR10 assembly.

### Simulations

To demonstrate the accuracy of our pipeline that produces copy number estimates derived from 12-mer abundances in sequencing reads, we performed 1,000 simulations wherein directed copy number changes were introduced into the reference genome assembly, Illumina sequencing reads were simulated from the genome with introduced copy number variation, and the linear relationship between the copy number estimates produced by our pipeline (based on 12-mer abundances in sequencing reads) was compared to the actual relative copy number of a random 1kb sequence for -1 to +10 copies. For each simulation, we first simulated background polymorphism by generating 100 random copy number variations ranging from 100 to 10,000 bp with a max copy number of 10 and copy number variation gain loss ratio of 1 in the *A. thaliana* reference genome using simulG v1.0.0 (Yue and Liti, 2019). We then selected a random 1 kb sequence and simulated deletion (-1 copy) to +10 tandem insertions copies of the selected sequence using simulG. Single-end 100 bp Illumina HiSeq 2000 reads at 5X coverage were simulated from each genome with directed copy number change with ART_Illumina v2.5.8 (Huang et al., 2012). Canonical 12-mers were counted in the simulated reads using jellyfish v2.2.9 (Marçais and Kingsford, 2011) and copy number estimates were produced using the last step of the pipeline depicted in Fig. S4. Linear models were fitted in R (R Core Team, 2013).

### Genome content profiles

Sequencing reads from 1,319 previously sequenced accessions of *Arabidopsis thaliana* (studies PRJNA273563 (1001 Genomes Consortium, 2016), PRJEB19780 (Durvasula et al., 2017), and SRP062811 (Zou et al., 2017)) were downloaded from the European Nucleotide Archive. The reads were adapter and quality trimmed with Trimmomatic 0.36 (Bolger et al., 2014) to remove sequencing adapters, low quality sequence on read ends (quality < 20), low quality sequence in sliding windows (clip at quality < 20 in 4 bp sliding window), the first 20 base pairs of each read, and reads less than 36 base pairs long. A custom adapter file containing all Illumina single end (SE) or paired end (PE) adapter sequences was used to trim adapters from SE or PE reads, respectively. Trimmed reads were then error corrected with BayesHammer (Nikolenko et al., 2013) as implemented in SPAdes 3.15.0 (Bankevich et al., 2012) using the default parameters. Error-corrected reads were then mapped to both the reference genome and the organellar genomes (Arabidopsis Genome Initiative, 2000) separately with bwa-mem 0.7.17 (Li, 2013; Li and Durbin, 2009). Alignment statistics were calculated with samtools 1.9 (Li et al., 2009). To reduce contamination from reads originating from the organellar genomes, reads that did not map to the organellar genomes were extracted from the alignments with bedtools 2.27.0 (Quinlan and Hall, 2010). Canonical 12 mers (i.e. sequences and reverse complements were binned together) were counted in the unaligned reads using jellyfish 2.2.9 (Marçais and Kingsford, 2011).

To remove samples that likely contained contamination, samples in which less than 90% of processed reads mapped to the reference genome were excluded from further analysis. Samples with coverage less than 1X based on coverage estimates produced using the sum of raw K-mer abundances found in the library divided by the approximate size of the genome (150 Mbp) were also subsequently disqualified. To remove libraries with extremes of GC content, which are likely due to PCR-bias during library preparation (Aird et al., 2011), samples with extreme GC content were filtered according to the 1.5 IQR rule.

To enable the comparison of 12-mer profiles between accessions, 12-mer counts for each sample were normalized according to a two step process. First, to correct for GC content bias due to variable library preparation across samples, raw 12-mer counts were transformed such that the proportion of counts that fell into GC bins defined by the GC content of each 12-mer sequence was constant across samples. As a “reference sample” for the transformation, the proportion of counts attributed to each GC bin was calculated using the median count for each 12-mer across accessions. For each sample, the proportion of counts attributed to each GC bin was tabulated and a weight was calculated as the ratio of the proportion of the “reference sample” counts in each GC bin to the proportion of the sample counts in each GC bin. Sample counts were then transformed by multiplying the raw counts by the corresponding weights of each GC bin. Samples with extreme Spearman’s correlation between GC-normalized 12-mer count and 12-mer GC content were also filtered according to the 1.5 IQR rule.

To normalize for coverage differences, GC transformed counts were normalized by quantile-quantile normalization using limma v. 3.44.3 R package (Ritchie et al., 2015). To reduce the impact of technical biases on 12-mer profiles, we regressed out the effect of the sequencing center by fitting a linear model.

Finally, near-isogenic accessions were filtered from the dataset. Genetic distance was calculated using published SNP genotypes (1001 Genomes Consortium, 2016) with plink 1.9 (Chang et al., 2015) and the distance matrix was hierarchically clustered and static tree cut at a height of 100,000 in R (R Core Team, 2013). For each cluster defined by the tree cut, one sample was randomly selected to be retained for further analysis and the remaining samples were filtered.

### Sequence abundance estimation with genome content profiles

The abundance of annotated sequences were estimated in each genome content profile by extracting the overlapping 12-mers that compose the feature and calculating the median count of these representative 12-mers (Fig. S4). This value was then transformed to an abundance estimate according to the following exponential equation: 2^x^, where x is the median count of representative 12-mers. For all sequences besides the genomic windows, the abundance estimates were normalized by the median abundance estimate for complete single copy BUSCO genes within each sample. The *A. thaliana* Repbase 23.10 library (Bao et al., 2015) was used as the library of sequences for which copy number was estimated and supplemented with a 5S rDNA sequence from GenBank (GenBank M65137.1), a 45S rDNA sequence (Rabanal et al., 2017), and consensus sequences of each of the six 180 bp centromere satellite variants described by (Maheshwari et al., 2017). Consensus sequences for each of the centromere satellite variant clusters was determined by further clustering sequences from each cluster with vsearch 2.9.1 (Rognes et al., 2016) with a 75% identity threshold. The centroid that represented the greatest number of sequences per cluster was then used as the representative sequence for that cluster.

### Sequence abundance variability across accessions and groups

The standardized range (sR), defined as the range divided by the median of sequence abundance across accessions, was used to estimate sequence abundance variability.

To assess differences in repeat abundance across populations, we first calculated z-scores for the normalized abundance estimates. We then applied hierarchical clustering to the resulting matrix, clustering both individuals (columns) and sequences (rows). To test the clustering of individuals within subpopulations or sequences by class, we used a permutation approach. Specifically, we compared the within-cluster sum of squares of observation ranks to 10,000 random shufflings of the categorical labels to generate an empirical p-value for each observation.

To identify sequences with enrichment for copy number increases or decreases per subpopulation, we asked whether there was an enrichment of individuals within a subpopulation with extreme scaled abundance estimates suggestive of copy number increase (Z > 3) or copy number decrease (Z < -3) compared to all other samples. Enrichment was calculated using Fisher’s Exact test as implemented in the R stats package (R Core Team, 2023). P values were corrected for multiple testing using Bonferroni’s method, with the number of tests corresponding to the number of groups for each sequence.

### Variant filtering and genome-wide association

Published variant calls from the 1001 Genomes Project (1001 Genomes Consortium, 2016) were filtered with bcftools 1.8 (Li et al., 2009) and plink 1.9 (Chang et al., 2015) to retain biallelic sites called in 95% of individuals with a minimum minor allele frequency of 0.01 and minimum quality of 30.

Sequence abundance was treated as a molecular phenotype and mapped using genome-wide association with a mixed-linear model with GEMMA 0.98 (Zhou and Stephens, 2012). Only sequences with a sR greater than the 99th percentile of sR across BUSCO genes were included in the analysis. A centered relatedness matrix was used to correct for population structure. K-mer sequence abundance estimates were log_2_-transformed prior to analysis. Bonferroni threshold (α/*N*, *w*ℎ*ere N is t*ℎ*e number of variants*) was used to determine significant variants for individual studies.

To determine if significant SNPs were enriched in the pericentromere and centromere, one-sided Fisher’s Exact tests were applied to 2 × 2 contingency tables describing whether SNPs were significant or not or found within the pericentromere and centromere or chromosome arms. Only GWAS with 10 or more significant SNPs were included in this analysis. Coordinates for the pericentromeres and centromeres in the reference genome are attributed to (Underwood et al., 2018). A Bonferroni-adjusted threshold was used to determine significance for these tests.

GWA with enrichment for significant SNPs in the pericentromere and centromere were considered to be localized to a specific centromere if they had at least half of their significant SNPs present within the centromere.

The effect of an allele on sequence abundance was calculated by taking the harmonic mean of sequence abundance across individuals possessing the allele and comparing the value to the respective mean for individuals with the alternative allele.

### Meta-analyses of genome-wide association of sequence abundance variation

Meta-GWAS analysis was performed by combining p-values from genome-wide association likelihood ratio tests using Fisher’s method per the following equation: 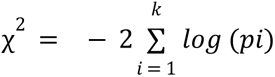 (Evangelou and Ioannidis, 2013) where *k* is the number of studies included in the meta-analysis and *p_i_* is the p-value for a variant in study *i.* Individual studies with excessive numbers of significant SNPs and studies in which the K-mer based sequence abundance estimates were strongly correlated (Spearman’s *ρ* > 0.60) with one or more other sequences, were excluded from analysis.

To correct for test statistic inflation across studies included in the meta-analyses, the genomic inflation factor, λ, was calculated and used to correct the chi-squared values. λ was calculated as follows: λ = *median* (χ^2^ *observed*) / *median* (χ^2^ *expected*) where the median expected χ^2^ value is the value at the 0.5 / k quantile of observed values. Focal variants were considered to be the set of SNPs with the top 0.1% of χ^2^ value per meta-analysis.

### Gene ontology enrichment analysis

Gene ontology enrichment analysis was done with topGO 2.46.0 (Alexa and Rahnenfuhrer, 2019) using the “weight01” algorithm and Fisher’s exact test. Genes of interest were considered to be genes within 10 kb of tag SNPs under each meta-GWA peak. Genes were considered to be significantly enriched if p < 0.05.

### Evolutionary analyses

Allele frequencies were calculated using plink 1.9 (Chang et al., 2015). Alleles were assigned ancestral or derived state based on an alignment of the TAIR10 genome assembly to the *Arabidopsis lyrata* v.1 assembly (Hu et al., 2011) using minimap2 v. 2.17 (Li, 2018) using the “asm10” parameter. Only sites with segregating ancestral and derived alleles in *A. thaliana* were considered for further analysis. Variants were classified as nonsynonymous or synonymous based on annotating the 1001 Genomes VCF using the Araport11 annotation (Cheng et al., 2017) and Variant Effect Predictor v. 103.1 (McLaren et al., 2016). Associated variants from AraGWAS (Togninalli et al., 2020, 2018) were accessed on 2021-09-01 and consisted of 44,680 significant associations across 462 studies.

Sequences with copy number change were identified by testing whether the median sequence abundance per group was different from the median abundance in Africa, the presumed progenitor to all other groups, using Wilcoxon-Mann Whitney U test in R (R Core Team, 2013). Bonferroni-adjusted alpha of 0.05 was used to determine significance.

## Acknowledgments

We thank Danelle K. Seymour, Keely E. Brown, Jacob B. Landis, Jason E. Stajich, and Zhenyu (Arthur) Jia for their helpful comments and suggestions that improved this work. All computational analyses were performed using resources at the High Performance Computing Center at the University of California, Riverside, supported by NSF grants MRI-2215705 and MRI-1429826, and NIH grant 1S10OD016290-01A1. D.K. is supported by NSF CAREER grant IOS-2046256.

## Data Availability Statement

A user-friendly generalizable version of the K-mer pipeline is available at GitHub at https://github.com/cjfiscus/Kmer-it. The 12-mer based sequence abundance estimates along with an archived version of the K-mer pipeline that was used to generate the data for this manuscript are available from figshare at https://figshare.com/articles/dataset/Resources_for_Fiscus_CJ_Koenig_D_The_genetic_control_of_rapid_genome_content_divergence_in_Arabidopsis_thaliana_bioRxiv_2025_p_2025_06_11_659220_doi_10_1101_2025_06_11_659220_/29422016. All other data and methods required to reproduce this work are described within the manuscript or the supplementary material.

