## Supporting Information for "The genetic control of rapid genome content divergence in *Arabidopsis thaliana*"

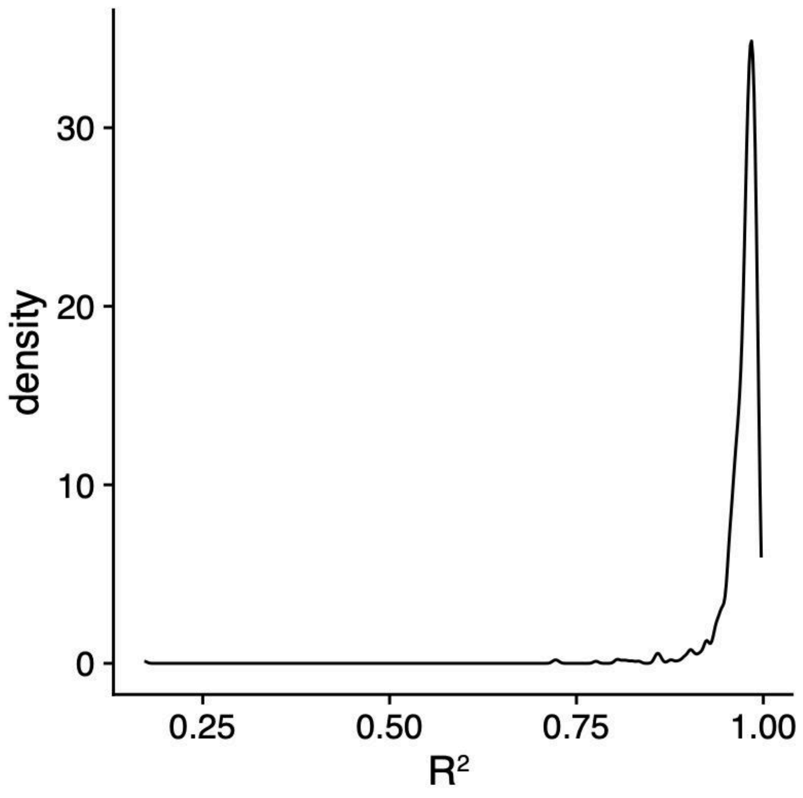


**Figure S1. Distribution of $R^2$ values from linear models to predict simulated copy number variation using 12-mer abundance**

Each model compares the actual and 12-mer estimated copy numbers for a single 1 kb sequence selected from the reference genome, with simulated copy number from -1 (deletion) to 10 copies.

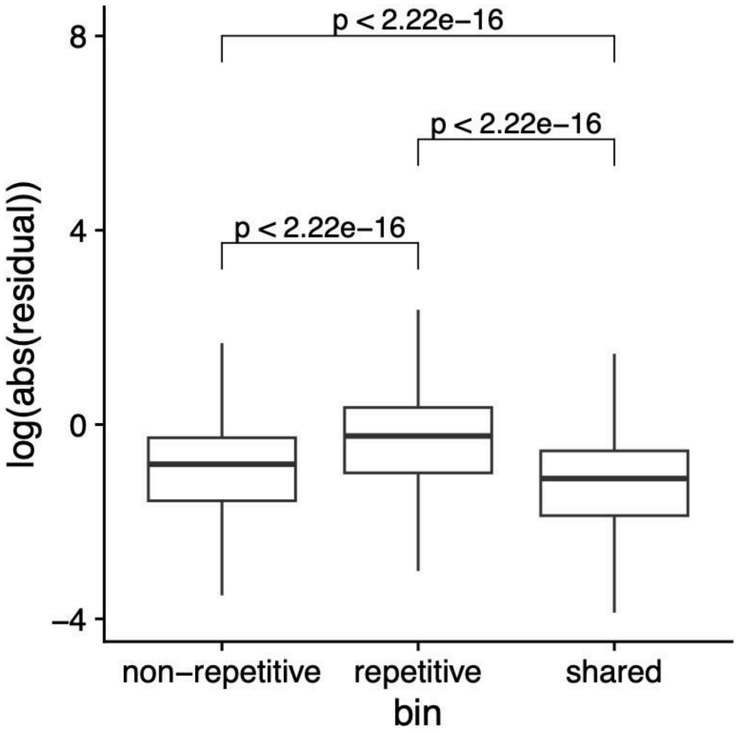


**Figure S2. Box plots of residuals from linear regression between 12-mer abundances in the TAIR10 reference genome and high-throughput sequencing reads**

P values were calculated using Wilcoxon rank-sum tests comparing median residuals per group. Outliers are not shown.

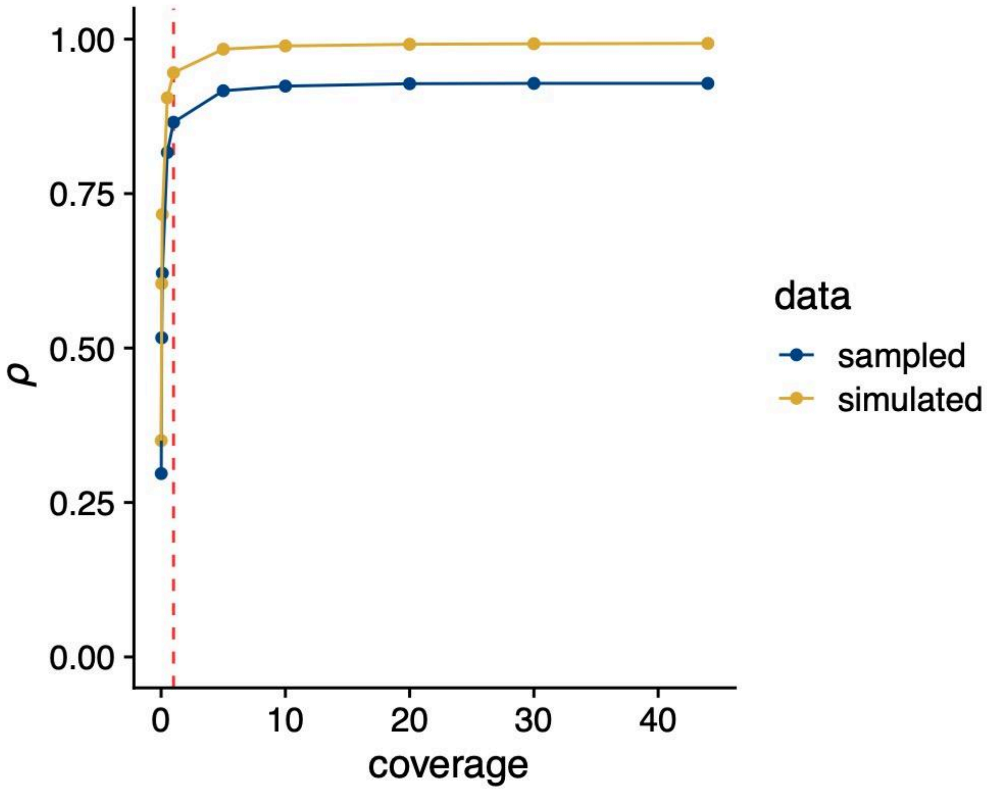


**Figure S3. Correlations between 12-mer frequencies in sampled or simulated high-throughput sequencing reads and the TAIR10 reference genome**

The dotted red line denotes 1X coverage, which was used as the threshold for coverage-based filtering in the pipeline. Sampled reads were generated by downsampling *Col-0* sequencing reads from the 1001 Genomes Project to each target coverage. Simulated reads were generated from the TAIR10 reference genome.

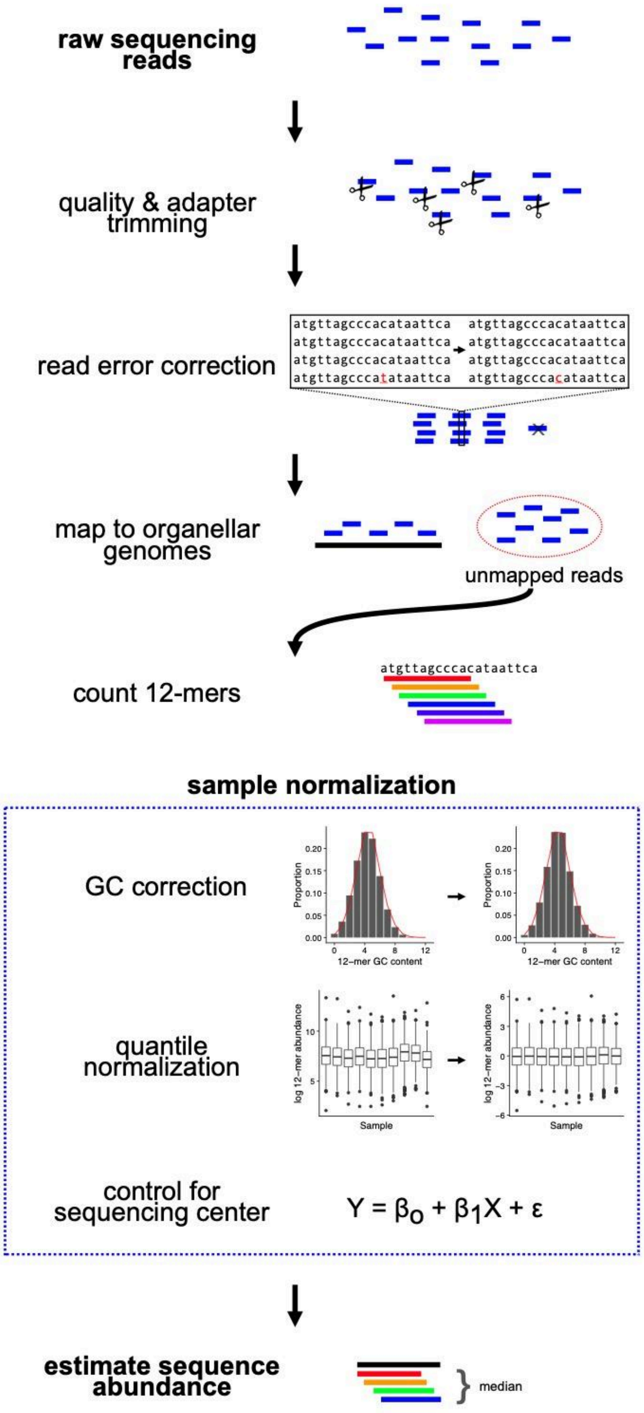


**Figure S4. Pipeline for generating GCPs and estimating sequence copy number from 12-mers**

The pipeline takes raw sequencing reads as input, which are trimmed for quality and adapter sequences using Trimmomatic. The trimmed reads are then error-corrected with SPAdes and mapped to the organellar genomes using BWA-MEM to remove organellar reads. The remaining unmapped reads, presumed to originate from the nuclear genome, are used for 12-mer counting with Jellyfish. After 12-mers have been counted for all samples, counts are normalized first for GC bias and then for sequencing coverage, followed by linear model correction to remove sequencing center effects. The resulting GCPs are then used to estimate the copy number of focal sequences by taking the median normalized 12-mer count across all overlapping 12-mers comprising the focal sequence.

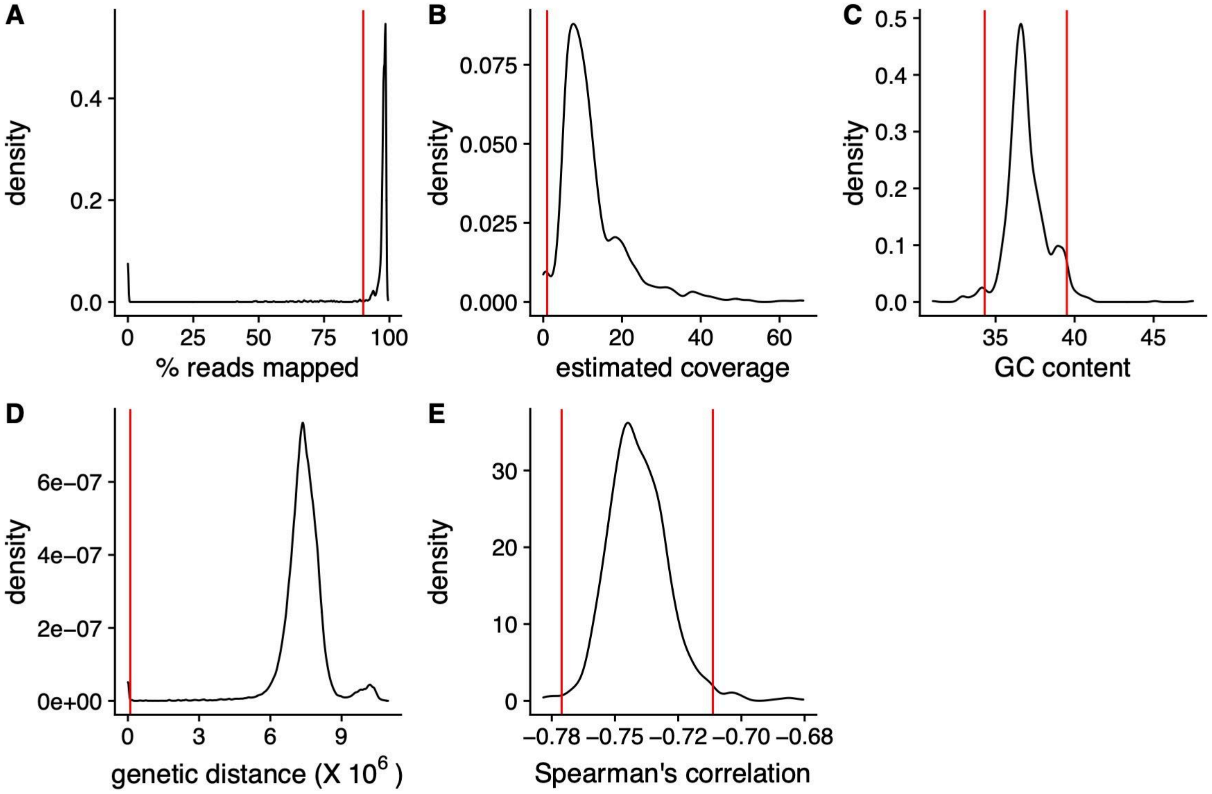


**Figure S5. Quality control filters for the 12-mer counting pipeline**

Distributions of (A) the proportion of reads mapped to TAIR10 genome per sequencing run, (B) sequencing coverage estimated from 12-mer counts, (C) GC content per sequencing run, (D) pairwise genetic distance (allele count) and (E) Spearman's correlation between GC content and 12-mer abundance. Vertical red lines indicate hard filtering thresholds. In panels A, B, and D, samples with values below the threshold were filtered. In panels C and E, samples with values outside the range denoted by the vertical lines were filtered.

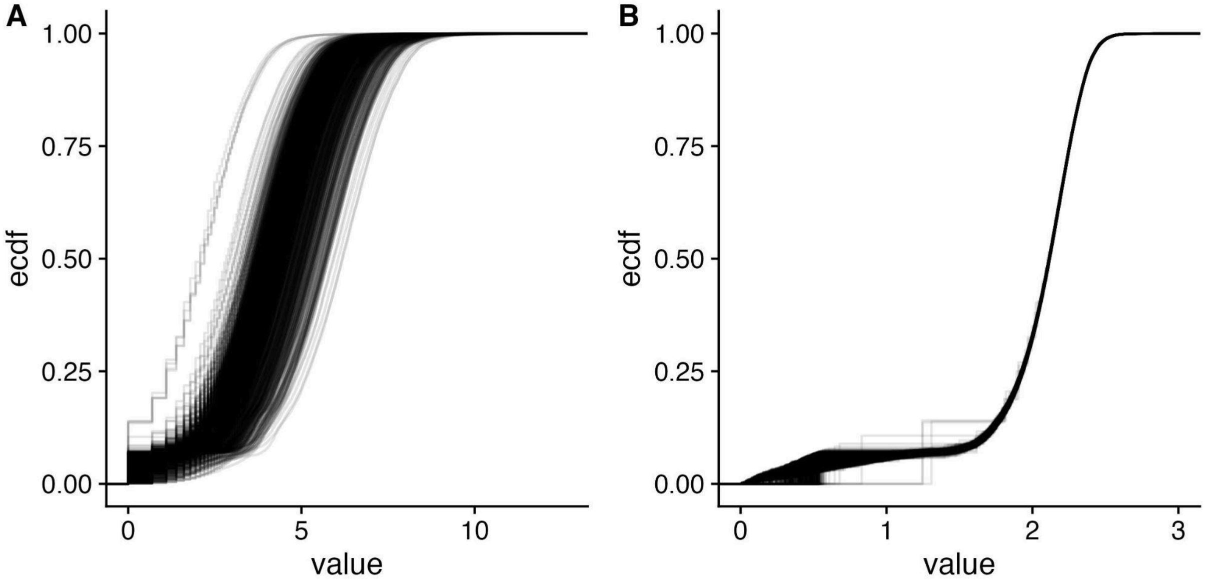


**Figure S6. Effect of frequency normalization on 12-mer abundance distributions**

Empirical cumulative distribution functions of 12-mer abundances for 10,000 random K-mers before (A) and after (B) quantile-quantile (QQ) normalization.

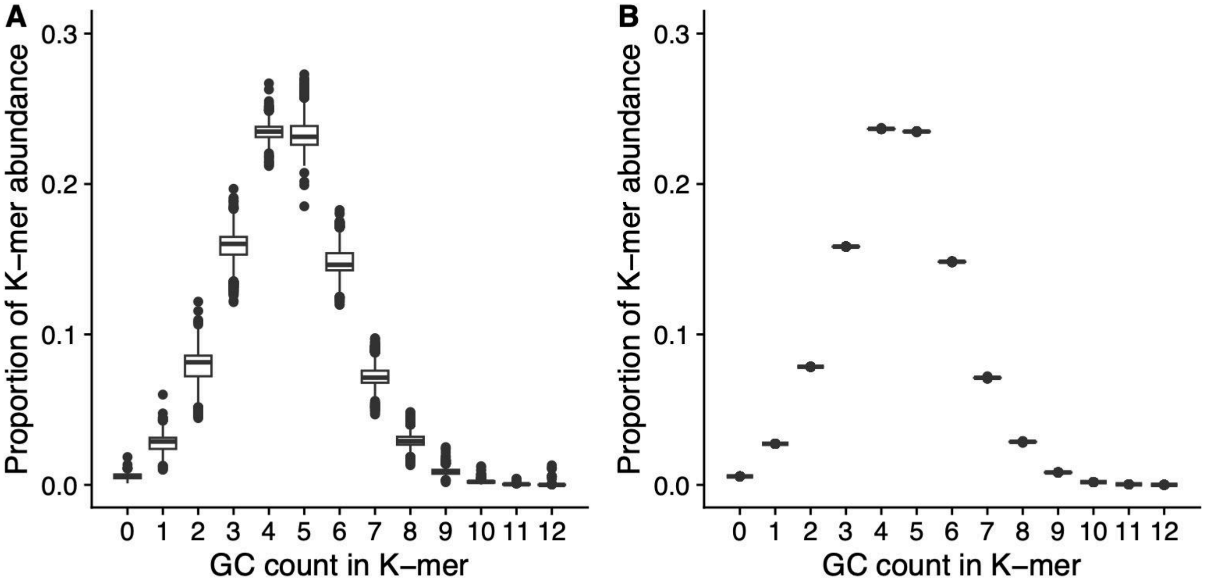


**Figure S7. Effect of GC normalization on the GC content dependence of 12-mer profiles**

Tukey boxplots of 12-mer abundances binned by 12-mer GC content before (A) and after (B) GC normalization.

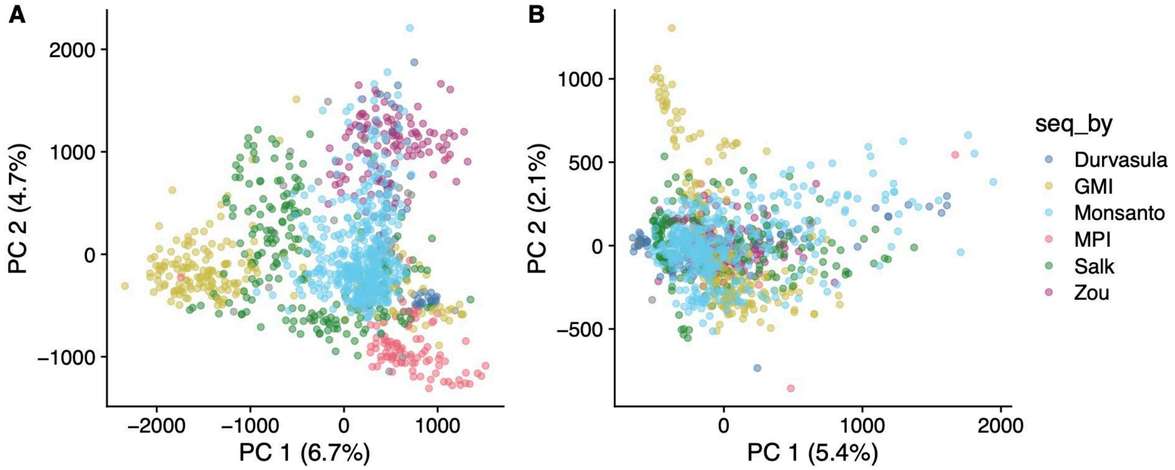


**Figure S8. Effect of correcting for sequencing center on principal component analysis of 12-mer profiles**

(A) Principal component analysis of 12-mer profiles before correction for sequencing center, with points colored by sequencing center. (B) Principal component analysis of 12-mer profiles after correction for sequencing center.

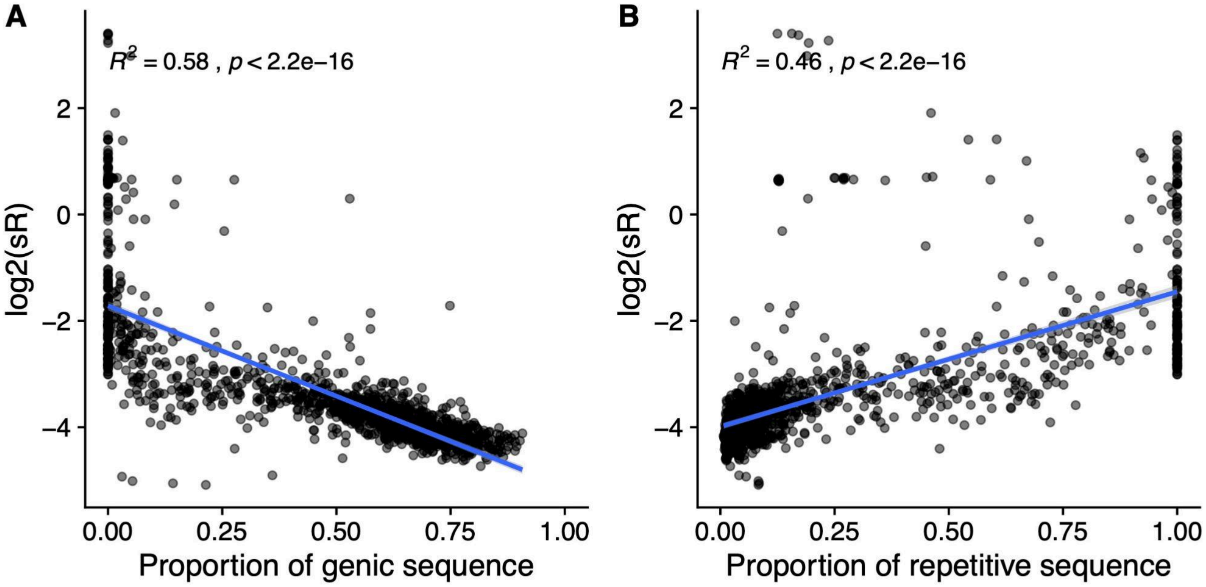


**Figure S9. Relationship between sR and reference genome sequencing composition in 100 kb windows.**

Abundance variability (log-standardized range; sR) per 100 kb window as a function of proportion of (A) genic sequence and (B) repetitive sequence. Blue lines indicate linear model fits.

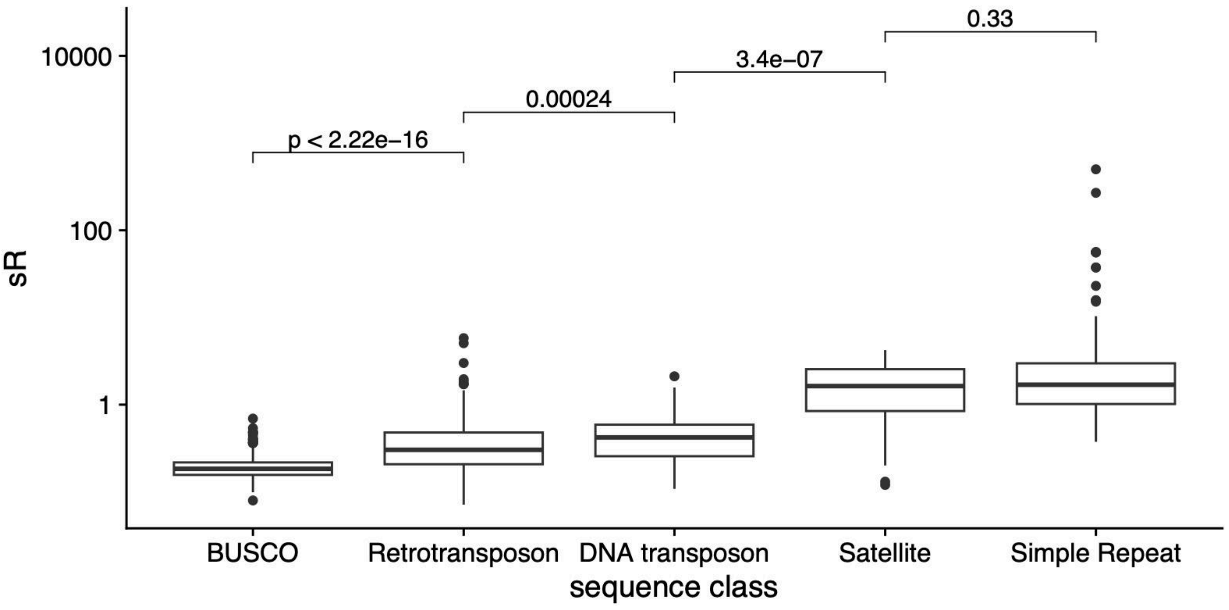


**Figure S10. Abundance variability of sequences by class**

Tukey boxplots showing sequence abundance variability measured as the standardized range (sR). P values were calculated from Wilcoxon-Mann Whitney U tests.

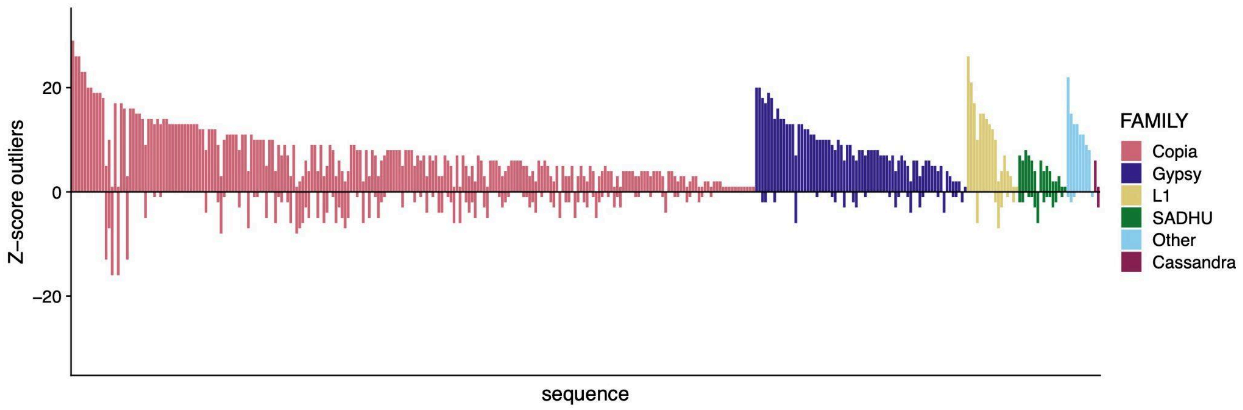


**Figure S11. Abundance outliers for retrotransposons by superfamily**

Each bar represents a retrotransposon family, and bar height indicates the number of accessions showing increased (positive values) or decreased (negative values) copy number relative to the population mean. Colors indicate superfamily.

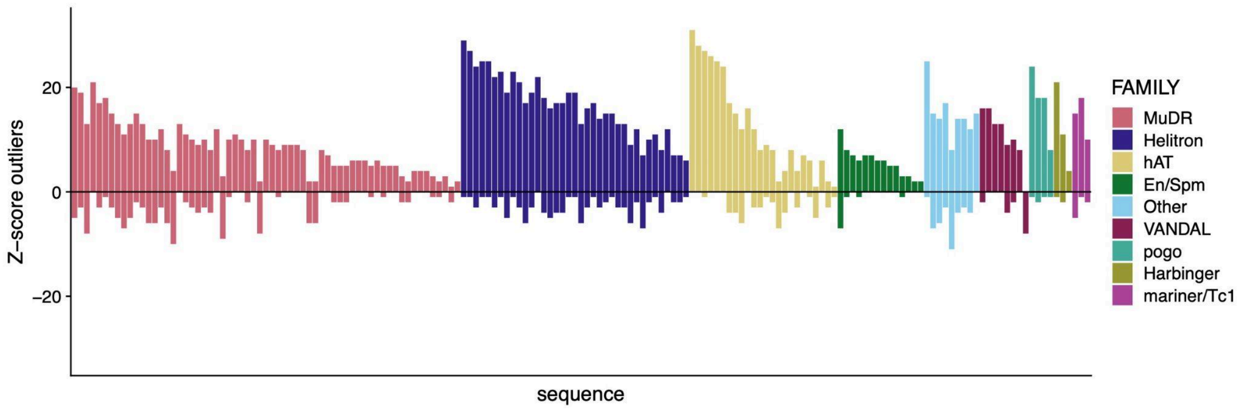


**Figure S12. Abundance outliers for DNA transposons by superfamily**

Each bar represents a DNA transposon family, and bar height indicates the number of accessions showing increased (positive values) or decreased (negative values) copy number relative to the population mean. Colors indicate superfamily.

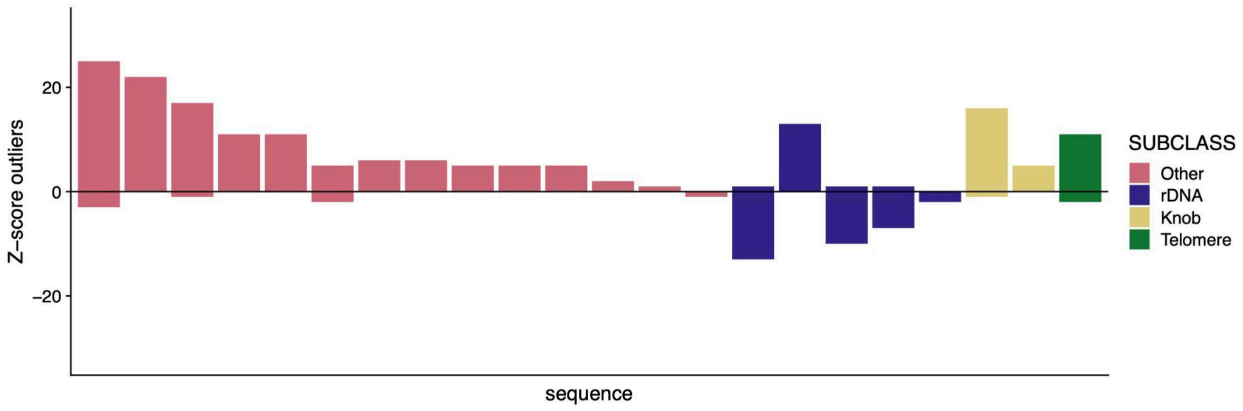


**Figure S13. Abundance outliers for satellites by subclass**

Each bar represents a representative satellite sequence, and bar height indicates the number of accessions showing increased (positive values) or decreased (negative values) copy number relative to the population mean. Colors indicate satellite subclass.

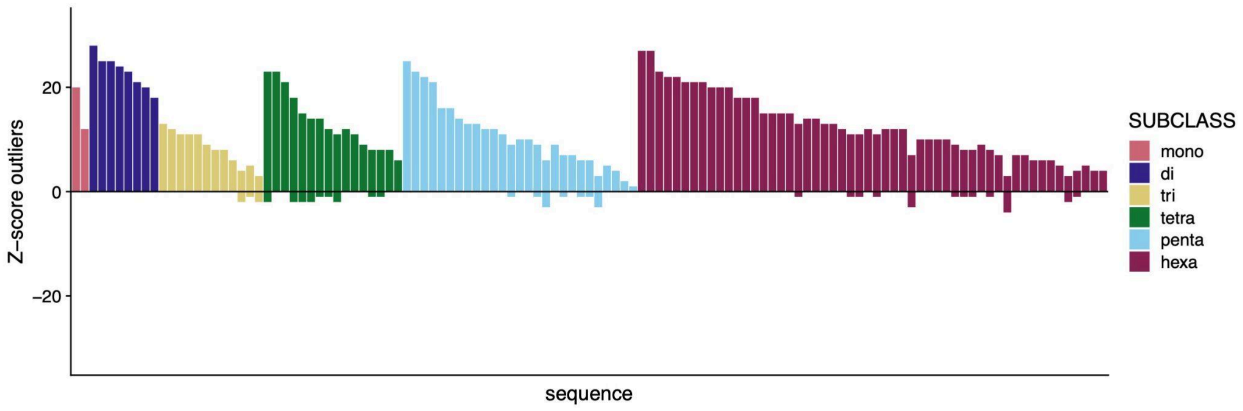


**Figure S14. Abundance outliers for simple repeats by subclass**

Each bar represents a representative simple repeat sequence, and bar height indicates the number of accessions showing increased (positive values) or decreased (negative values) copy number relative to the population mean. Colors indicate simple repeat subclass.

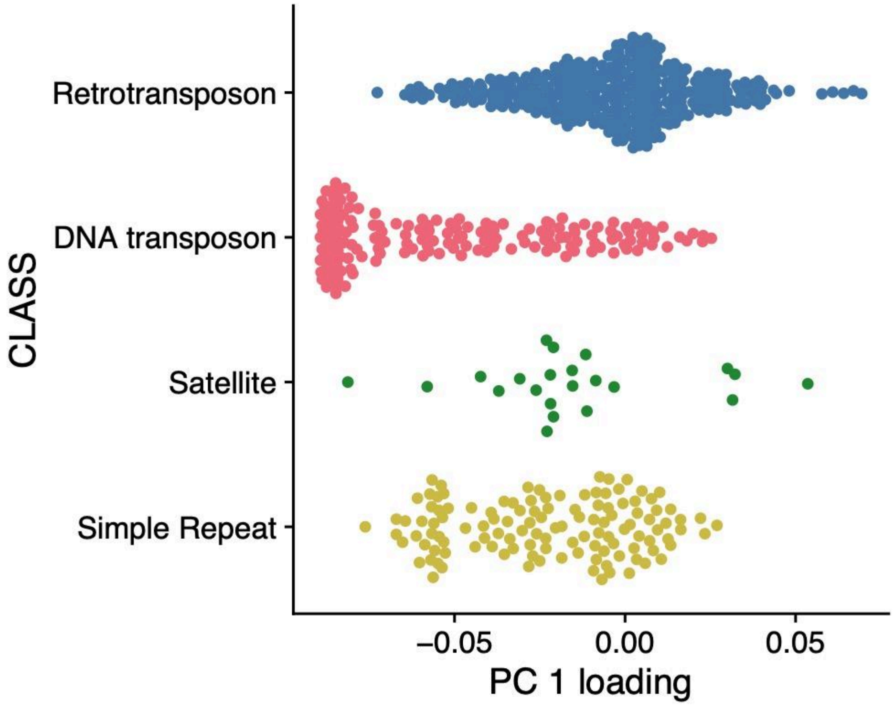


**Figure S15. Distributions of PC 1 loadings by repeat class**

Each point represents the PC1 loading of a repetitive sequence, with points jittered to show the distribution within each class.

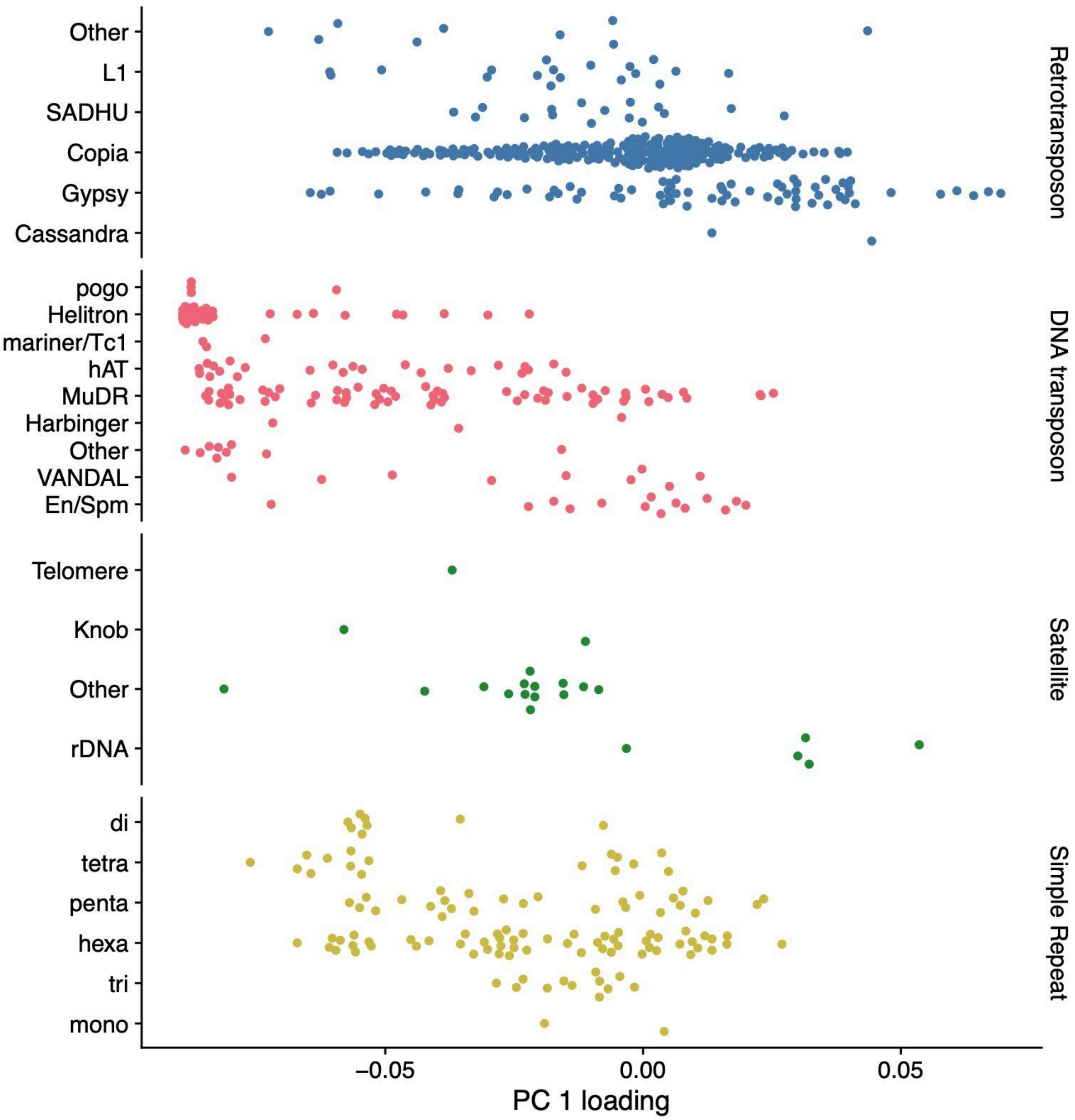


**Figure S16. Distributions of PC 1 loadings by repeat subclass**

Each point represents the PC1 loading of a repetitive sequence, with points jittered to show the distribution within each class. This is the same data shown in Figure S15 but by superfamily or subclass.

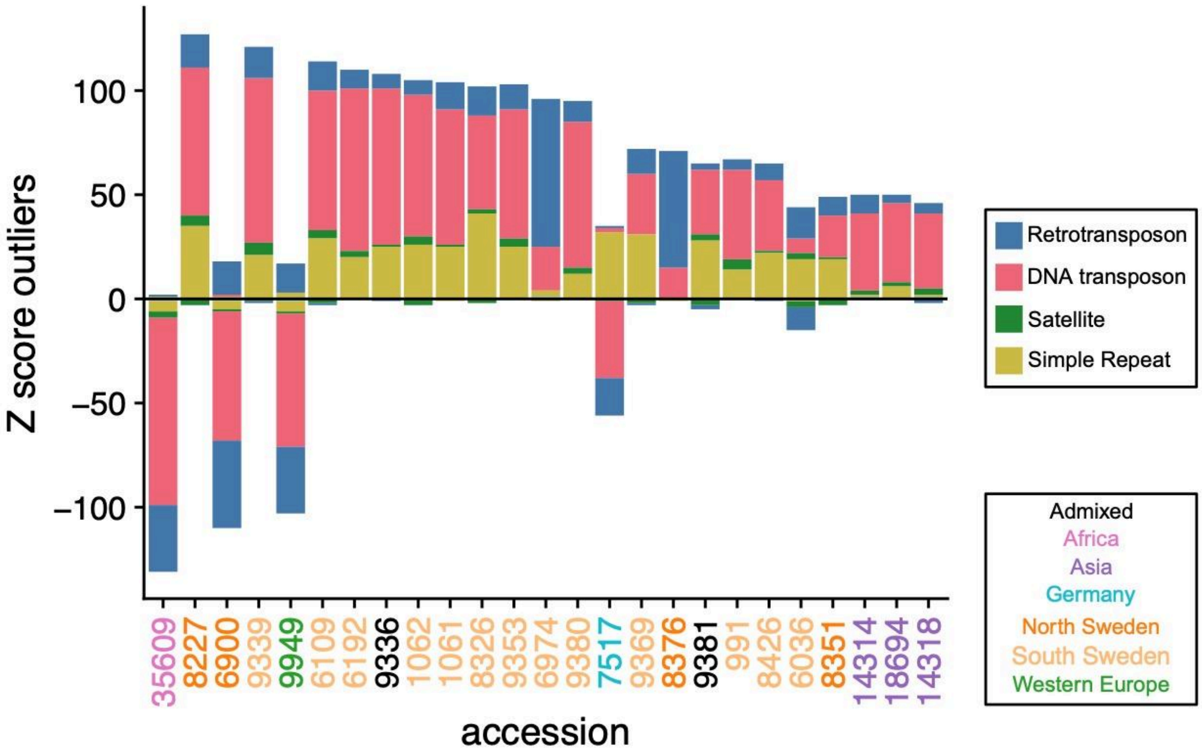


**Figure S17. Frequency of Z score outliers by sequence class for the top 25 most divergent GCPs**

Each column represents an accession, with labels colored by admixture group. Bar height indicates the number of sequences showing increased (positive values) or decreased (negative values) copy number relative to the population mean. Bars are filled according to repetitive sequence class.

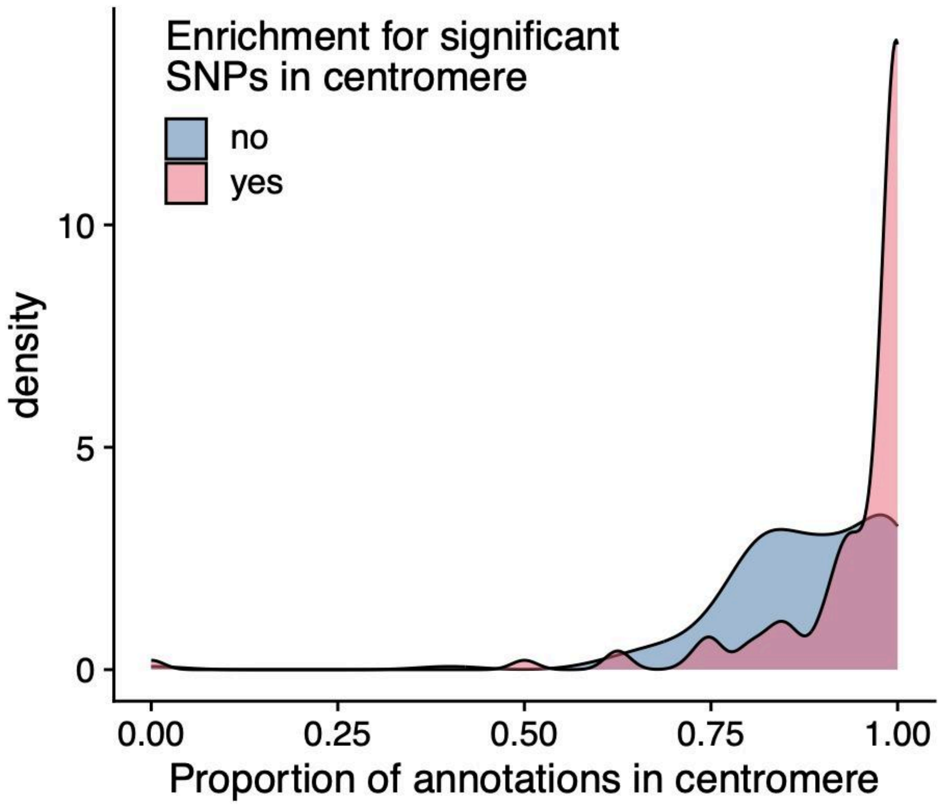


**Figure S18. Sequences with significant GWAS SNPs in the pericentromere and centromere are more likely to occur in centromeric regions**

For each GWAS with at least 10 hits, we calculated the proportion of sequence annotations located in the pericentromere and centromere, and compared distributions based on whether significant SNPs from the GWAS were enriched in these regions.

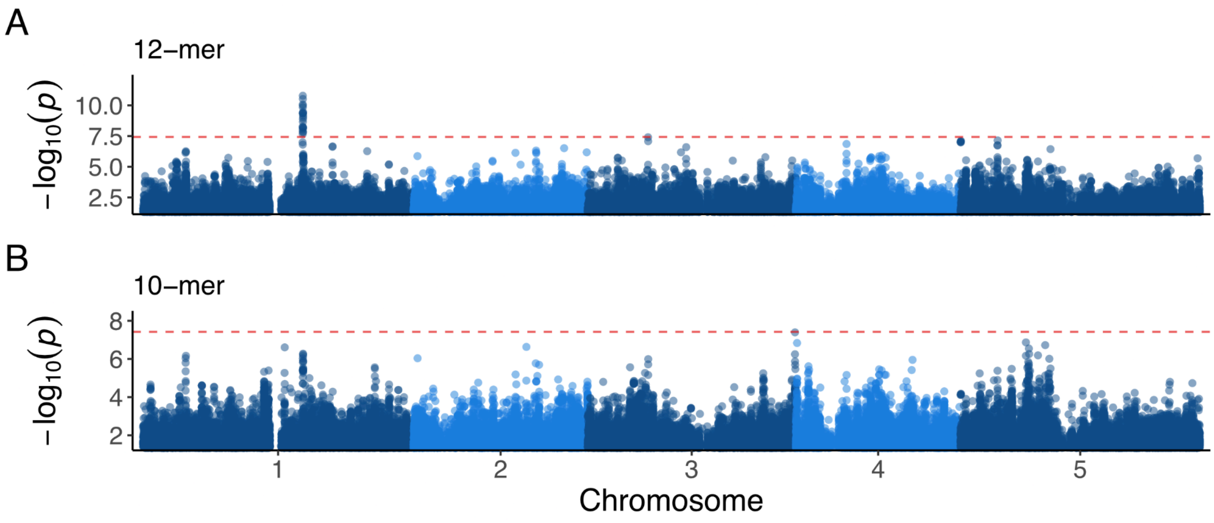


**Figure S19. 12-mer and 10-mer GWAS for ATCOPIA78_I**

Manhattan plots showing GWAS results for the internal sequence of *ONSEN* using 12-mer or 10-mer based GCPs. The dotted red line indicates the Bonferroni-adjusted significance threshold ($P < 0.05$).

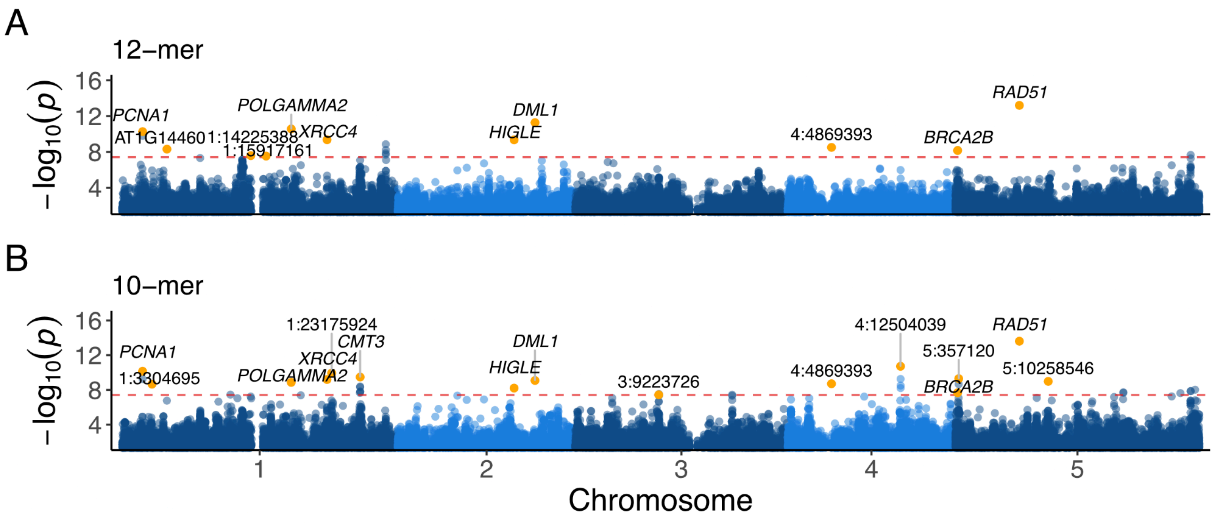


**Figure S20. 12-mer and 10-mer GWAS for ATHATN2**

Manhattan plots showing GWAS results for the representative sequence of ATHATN2 using 12-mer or 10-mer based GCPs. Meta-GWAS candidate SNPs are highlighted in orange and labeled with either the associated candidate gene or top SNP. The dotted red line indicates the Bonferroni-adjusted significance threshold (P < 0.05).

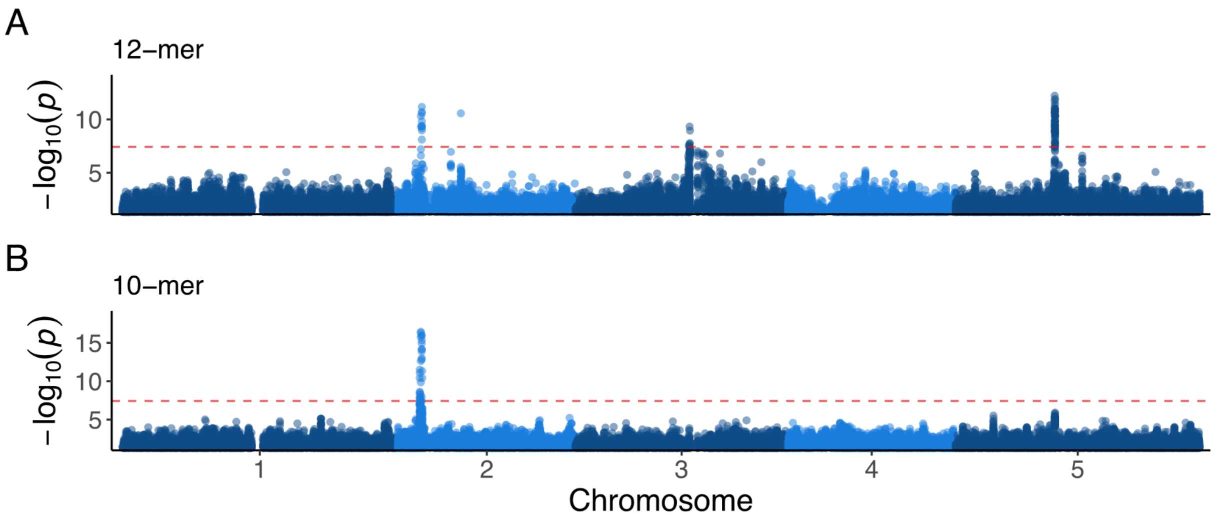


**Figure S21. 12-mer and 10-mer GWAS for VANDAL1**

Manhattan plots showing GWAS results for the representative sequence of VANDAL1 using 12-mer or 10-mer based GCPs. The dotted red line indicates the Bonferroni-adjusted significance threshold (P < 0.05).
